# Discordance between phylogenomic datasets in aphids: who is telling the truth?

**DOI:** 10.1101/2024.04.12.589189

**Authors:** Emmanuelle Jousselin, Armelle Coeur d’acier, Anne-Laure Clamens, Maxime Galan, Corinne Cruaud, Valérie Barbe, Alejandro Manzano-Marin

## Abstract

Aphids (Hemiptera: Aphididae) are intensively studied due to their significance as pests and their captivating biological traits. Despite this considerable research interest, the evolutionary history of this insect family is poorly understood. Recent phylogenomic analyses have produced conflicting topologies, particularly at deep nodes, complicating our understanding of aphid trait evolution. In this work, we aimed to produce new data to unravel the backbone phylogeny of aphids. We sequenced partial and whole mitochondrial genomes from 87 species that were added to 31 published mitochondria. We additionally sequenced 42 nuclear loci across 95 aphid species and sourced 146 genes from 12 new and 61 published genomes from the primary aphid obligate endosymbiont*, Buchnera aphidicola*. We obtain data from these three sources for a subset of 51 aphid species, facilitating a comparative analysis of their phylogenetic signals. Our analyses confirm the monophyly of subfamilies, validating current taxonomic classifications, except for Eriosomatinae and Calaphidinae. However, relationships between subfamilies remain contentious in both mitochondrial and nuclear phylogenies. The topologies obtained with *Buchnera* appear fully resolved but exhibit some discordance with host phylogenies at deep evolutionary scales and conflict with views on the evolution of aphid morphology. We discuss alternative hypotheses for these discrepancies. Finally, the paucity of phylogenetic information at deep phylogenetic scales may stem from an initial rapid radiation. Though challenging to establish, this scenario may inherently hinder resolution in aphid phylogenetics.

## Introduction

Aphids (Hemiptera: Aphididae) constitute a lineage of over 5,000 species of sap-feeding insects that include several significant agricultural, horticultural, and forestry pests (Blackman and Eastop 2000). This pest status has prompted numerous research programs aimed at a better understanding of their biology for the development of appropriate control methods. Beyond their notoriety as pest insects, aphids also exhibit variable biological characteristics, making them a prominent study system in evolutionary biology. For instance, their patterns of associations with host plants range from strict specialization towards a single host plant species to extreme polyphagy (with some aphid species capable of feeding on up to 100 plant families). Additionally, ten percent of all aphid species undergo seasonal alternation between two sets of host plants (Blackman and Eastop 2006, 2000), requiring the ability to locate and feed on plants belonging to different botanical families (Powell and Hardie 2001; Mackenzie and Dixon 1990; Jousselin *et al*. 2010). These features make aphids interesting systems for investigating how phytophagous insects adapt to their host plants and whether these adaptive processes drive speciation events (Shaposhnikov 1987, 1961; Heie 2004; Jousselin and Elias 2019). Aphids have also intensively been studied for their reproductive strategies undergoing cyclical parthenogenesis—alternating between sexual and asexual reproduction throughout their year-long life cycles. Both the ecological significance and genetic determinism of this biological character have attracted the interest of evolutionary biologists (Delmotte *et al*. 2001; Jaquiery *et al*. 2014). Aphids also stand out as a study system in bacterial endosymbiosis. Since Buchner’s pioneer work on endosymbiosis in invertebrates (Buchner 1965a), the study of their mutualistic association with their obligate nutritional symbiont, *Buchnera aphidicola*, has shed light on various aspects of the evolution of bacterial integration into eukaryotes. These aspects range from insect/symbiont cospeciation (Munson *et al*. 1991; Clark *et al*. 2000; Jousselin *et al*. 2009), metabolic cooperation (Feng *et al*. 2019; Wilson *et al*. 2010; Smith and Moran 2020) to degenerative evolution in endosymbiont genomes (Gil *et al*. 2002; Moran and Mira 2001; Toft and Fares 2009; Sabater-Muñoz *et al*. 2017). Many of the evolutionary biology studies cited above have been conducted on a few focal aphid species or groups. However, a thorough understanding of aphid biological evolution, requires knowledge of how aphid traits are distributed throughout the diversification of this lineage. Generating a highly accurate and robust aphid phylogenetic tree is a crucial step to that effect.

In addition to the reconstruction of the history of aphid biological traits, aphid phylogenetic history could help testing several macroevolutionary scenarios that have long been debated in the literature. The distribution patterns of aphid species through time and space is somewhat intriguing. The over 5,000 currently described aphid species (Favret, 2024) are organized into 23 subfamilies encompassing 510 genera. The first fossils identified as belonging to Aphididae date back to 160 My and the bulk of aphid fossil diversity is found in the Eocene (i.e. 56-33 My) (Heie 2004) and https://paleobiodb.org/ last accessed 14/12/2023). The 23 aphid subfamilies differ greatly in terms of number of species, with the most diverse subfamily, Aphidinae, gathering nearly half the diversity (with 2483 described species) while some species-poor subfamilies such as Phloeomyzinae for instance currently count no more than a single species. Differences in species diversity could be explained by the different ages of the subfamilies, with depauperate subfamilies being more recent and therefore having had little time to diversify and accumulate species. Alternatively, these variations in aphid species diversity could be caused by bursts of radiations and/or extinctions differentially affecting lineages of aphids. The latter scenario is clearly favored by taxonomists and paleontologists (Shaposhnikov 1981; Heie 2004; Heie 1987) and an early phylogenetic study (von Dohlen and Moran 2000). Another outstanding features of aphids, is their atypical geographic distribution pattern. They are highly diversified in the temperate region, and therefore show an inverse species diversity distribution to other insect groups (that are generally more diverse in the tropical region, Brown 2014). Several hypotheses have been put forward to explain this pattern. Those involve ecological factors such as a hyper specialization on a few host plants that might be maladapted to tropical environments (Dixon *et al*. 1987) or constraints linked to aphids historical biogeography (Shaposhnikov 1987). A thorough investigation of the scenarios underlying aphid species distribution through time and space necessitates a robust phylogenetic reconstruction. Successive burst of speciation and extinctions could be traced back, and historical biogeography scenarios could be untangled

Despite the wealth of scientific questions that necessitate a robust phylogenetic reconstruction, the aphid phylogenetic tree remains unresolved. The debate over the evolutionary relationships among aphid species groups actually predates the advent of molecular phylogenetics. This is well illustrated by the numerous taxonomic classifications that have been proposed over the years. The number and delimitation of aphid groups and their ranks (superfamily and subfamily) varied significantly among different authors. As Heie accurately pointed out, there were “*as many classifications as taxonomists*” (Heie 1980). Ilharco and Harten (1987) in their review of aphid taxonomic classifications listed four to 21 subfamilies grouped in one to ten families depending on authors and Wieczorek and Wojciechowski (2007) focusing solely on the group of taxa known as Drepanosiphine aphids (mainly Drepanosiphidae sensu Heie 1980) notes that this group included three to fourteen subfamilies depending on classifications. These taxonomic controversies were succinctly outlined in the introduction by Remaudière and Stroyan (1984), prompting the formulation of a classification featuring 20 subfamilies within a family named Aphididae. This classification was conceived with a functional perspective, wherein each subfamily grouped species (tribes, genera) perceived as homogenous by all taxonomists. Homogeneity, in this context, was defined by the exhibition of morphological characters distinctive enough to unequivocally differentiate and categorize aphids in a group (whether the groups was seen as a subfamily or a tribe by previous studies). There was thus no strong phylogenetic hypothesis nor evolutionary scenario underlying this classification. Attempts at higher-level classification, grounded in intuitive assessments of aphid general morphology rather than formal cladistic analyses (Quednau 2010), have then been sporadic. Remaudière’s (1984) classification gained popularity through the Aphid World Catalog (Remaudière and Remaudière 1997). Molecular studies have then often confirmed the monophyly of subfamilies (e.g. Chen *et al*. 2014; Chen *et al*. 2016; Choi *et al*. 2018; Wieczorek *et al*. 2017; Lee *et al*. 2022) validating Remaudière’s practical decision. Four additional subfamilies, each including very few species, have been delimited since then, and there are now 23 recognized aphid subfamilies (Favret 2024). But the relationships between these taxa remain unresolved (von Dohlen 2009; Podsiadlowski and Vilcinskas 2016). A few published molecular phylogenetic studies have focused on solving this puzzle. Among the most influential are the study of Novakova *et al*. (2013) using five DNA markers found on the primary symbiont genome of aphids, *Buchnera aphidicola*; the study of Chen *et al*. (2017) based on the whole mitogenomes from 33 aphid species and two recent phylogenomic investigations using respectively transcriptome data (Owen and Miller 2022) and ultra-conserved Elements (Hardy *et al*. 2022) each yielding thousands of nuclear markers (Fig. 1). These studies represent solid attempts at solving the backbone phylogeny of Aphididae in terms of taxa representation as they include several subfamilies and mostly subfamilies that encompass the bulk of aphid diversity. They also rely on various and sometimes numerous DNA markers. However, despite their endeavors, the resulting trees exhibit uncertainties in deep nodes. Many relationships between subfamilies lack robust support and depend on the phylogenetic inference methods employed. Additionally, relationships between subfamilies differed between studies with only a few deep nodes shared among all four trees (Fig. 1). Nevertheless, this comparison is inherently challenging due to the lack of overlap in taxa between studies. Consequently, discerning whether the incongruence in datasets arises from differences in taxa sampling, inference methods, or actual distinctions in phylogenetic signals based on DNA sources remains a complex task.

**Fig. 1:**
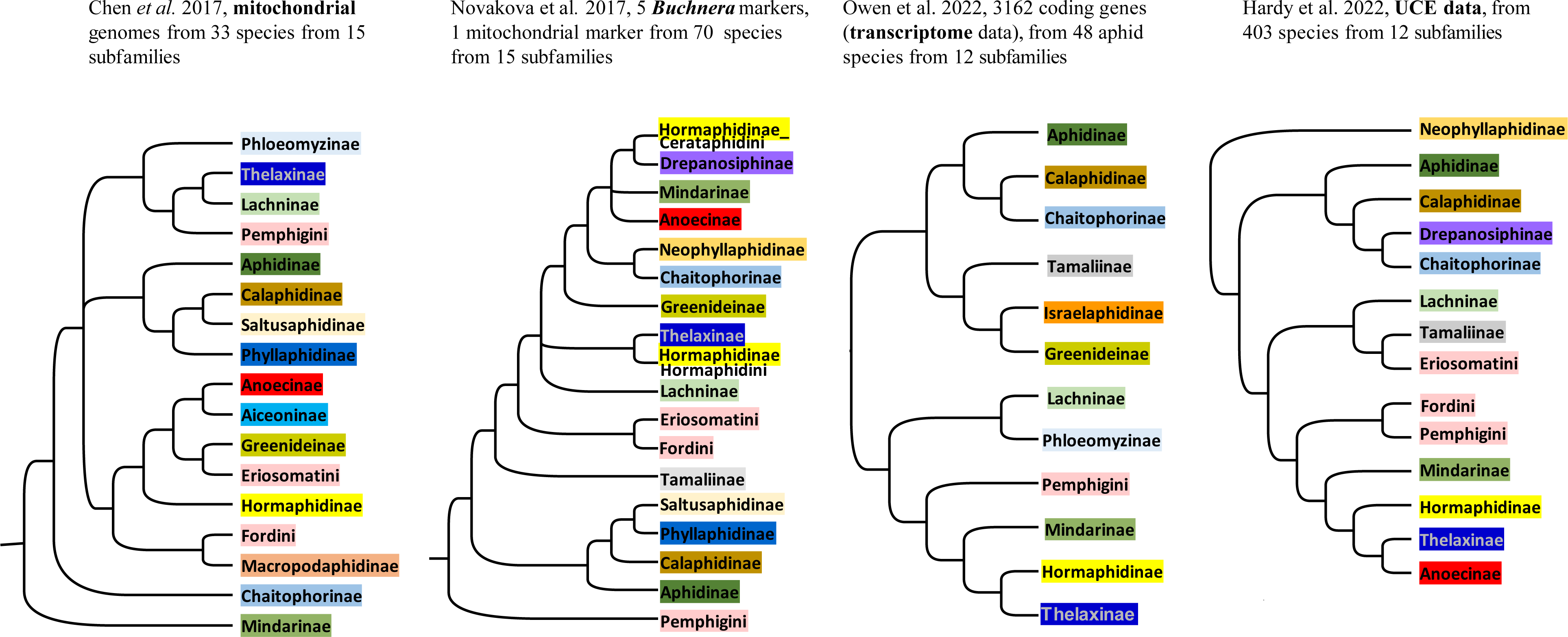
Alternative topologies found in previous phylogenetic investigations of aphids: 1) sketch of the Bayesian topology on Fig. 1 of Chen *et al*., 2017. 2) sketch of the Bayesian topology on Fig 2 of Novakova *et al*., 2013, 3) sketch of the ML topology on Fig. 1 of Owen *et al*. 2022, 4) sketch of the phylogeny obtained by Hardy *et al*. 2022.

Here, we aimed at obtaining data from three sources: the mitochondrial genome of aphids highly conserved nuclear genes from the aphid genome and genes from the aphid primary symbiont on a set of species representative of aphid diversity. This data was used to understand whether discrepancies between previously published phylogenies stem from uneven taxa representation and attempt solving aphid subfamilies relationships. First, we compiled a mitogenome dataset aiming to add subfamily representation to the study of Chen et al (2017) and expand representation of highly diverse subfamilies; to that effect we produced partial or whole mitogenomes that were combined to previously published mitochondria. We also generated multiple independent single-copy nuclear gene sequences, using the Fluidigm Access Array technology and high throughput sequencing. We then compared the phylogenetic information yielded by these two datasets with the phylogenetic history of *Buchnera aphidicola*. Those were obtained from *Buchnera* whole genome data mainly produced by Chong *et al*. (2019) and Manzano-Marín *et al*. (2023) to which we added 12 new draft genomes to cover additional subfamilies. We tackled how gene partitions obtained from mitochondrial, nuclear and endosymbiont data align with one another. We further assessed whether the relationships retrieved were congruent with previous views of the evolution of this well-known insect lineage and discuss potential sources of conflicts.

## Material and method

### Taxon sampling

Aphids samples used in this study were sourced from the CBGP INRAe aphid collection (DOI: 10.15454/D6XAKL), originally collected from 2006 to 2019 and preserved in 70% alcohol at 6°C. For each sample collected, a preliminary classification of individuals was carried out through observation under the microscope. Subsequently, to confirm the first visual morphological classification, some specimens were mounted on slides and formally identified by A. Coeur d’acier using mainly the keys of Blackman and Eastop (http://www.aphidsonworldsplants.info/). After morphological identification under the stereo-microscope, one to 10 specimens, representative of each sample, were randomly selected for DNA extractions and molecular analyses. A full list of samples newly sequenced for this study, voucher numbers as deposited in the CBGP INRAe aphid collection and sources from previously sequenced aphids are given in Table S1. Taxonomy follows (Favret 2024).

### Mitogenome data

In order to obtain full mitogenomes, we first retrieved reads from the Illumina sequencing of aphid endosymbionts from our previous studies (Manzano-Marín *et al*. 2020; Manzano-Marín *et al*. 2018; Manzano-Marín *et al*. 2023) and a dataset from Chong *et al*. (2019) downloaded from the NCBI Short Reads Archives. They generally yielded aphid mitogenomes with sufficient coverage to allow full assembly. DNA library construction and sequencing protocol are described in respective publications. For both these datasets, reads were trimmed and quality filtered using FASTX-Toolkit v0.0.14 (http://hannonlab.cshl.edu/fastx_toolkit/). Reads shorter than 75 base pairs (bp) were dropped. Additionally, PRINSEQ v0.20.4 was used to remove reads containing undefined nucleotides as well as those left without a pair after the filtering and clipping process. The resulting reads were assembled, using SPAdes v3.10.1 (Bankevich *et al*. 2012), with the options --only-assembler option and k-mer sizes of 33, 55, 77, 99. Contigs representing aphid mitogenomes were identified based on BlastX searches against a database containing available aphid mitogenomes in NCBI. The cox1 genes retrieved in our assemblies were systematically used in BlastN searches against our own curated barcode database (based on preparation of specimen vouchers) (Coeur d’acier *et al*. 2014) to confirm species identification and ensure that we did not have any cross-contamination or chimeric sequences. Mitogenomes generally assembled into a single contig. Through this process, 55 new aphid mitogenomes were assembled. Thirty-eight were obtained from endosymbiont sequence dataset from the INRAe-CBGP collection while 17 were obtained from the endosymbiont sequence dataset of Chong *et al*. (2019) (Fig S1, Table S1).

In order to cover additional subfamilies, we used PCR amplification of three overlapping mitochondrial DNA fragments (long fragments from about 4 kb to 8 kb) on several specimens. For each aphid sample, about ten to fifteen individuals from the same aphid colony were pooled together to obtain DNA extracts of over 10 ng/μL. For DNA extraction, we used the DNeasy Blood and Tissue Kit (Qiagen) following the manufacturer’s recommendations. DNA extractions were then normalized to 10 ng/ μL after Qubit fluorimeter quantification (Invitrogen). Long range PCR amplification and DNA library construction protocols from pooled amplicons and further sequencing are detailed in Supplementary Material (Text S1). Reads filtering, trimming, assembly and taxonomic assignments of contigs followed the same procedures as for the endosymbiont sequencing reads. All resulting contigs of the assemblies were identified as parts of an aphid mitogenome except for a few chimeric contigs or some contigs belonging to aphid parasitoid mitogenomes. Through this long-range PCR procedure, we obtained 28 partial and 4 complete mitochondrial genomes (Table S2). For each aphid sample, the contigs were scaffolded against the most closely related aphid mitogenomes available in NCBI or our own assemblies.

All mitogenomes generated were annotated using the MITOS Web server (Bernt *et al*. 2013) and annotations were curated manually using UGENE (10.1093/bioinformatics/bts091) and the Geneious 11.1.5 interface. In brief, we verified start-stop position of protein coding genes using annotated reference mitogenomes of Wang *et al*. (2013), we located and manually annotated a small number of tRNA genes that were missed using blastn searches. All complete and partial mitogenomes obtained in this study were submitted to NCBI using GB2sequin (Lehwark and Greiner 2019). Finally, we completed this dataset with 29 aphid mitogenomes and those of two outgroup species, *Daktulosphaira vitifoliae* and *Adelges tsugae*, available in NCBI. We chose sequences that were complete, preferentially obtained from aphid whole genome sequencing, in order to detect gene rearrangements if any. Altogether, we gathered 120 mitogenomes (Table S1).

The gene sequences were aligned using Mafft v7.2.2.1 (Katoh and Standley 2013). Alignments for protein-coding genes were corrected to respect translation frames when needed and were translated into amino acid sequences (AA). We produced two data matrices for further phylogenetic analyses: a full DNA matrix concatenating all 13 protein coding genes (tRNA sequences and ribosomal RNA genes were discarded from analyses as they provided little phylogenetic information or were too ambiguously aligned), and the corresponding AA matrix.

### Selection of single copy nuclear genes, high-throughput PCR on Fluidigm Access Array technology and high-throughput sequencing

#### Nuclear gene selection, primer design and validation

We first targeted nuclear markers that have been previously used in aphids phylogenetic investigations (Elongation factor and long-wave Opsin, Ortiz-Rivas and Martinez-Torres 2010) and then added single copy genes that have been used in scale insects phylogenetic investigation (Mullen *et al*. 2017). We further targeted single copy nuclear genes commonly used to solve deep nodes in insect phylogenies (within Lepidoptera Wahlberg and Wheat 2008; Wahlberg *et al*. 2016) and between insect orders (Wiegmann *et al*. 2009) but never tested on aphids. In order to define appropriate primers for aphids, these genes were searched for in aphid reference genomes available at the time of initiating this study: *Acyrthosiphon pisum* (Acyr 2.0), *Diuraphis noxia* (Dnoxia_1.0), *<u>R</u>hopalosiphum maidis* (ASM367621v3), *Eriosoma lanigerum* (WAA_JIC_1.0), *Sipha flava* (YSA_version1) and the outgroup *Daktulosphaira vitifoliae* (ASM2509136v1) using either available genome annotations or TblastN. For the later, query sequence used for blast searches was the pea aphid sequence of the targeted gene (using min Identity = 80). Genes that had blast hits against all available aphid genomes were then selected along for primer designs using Primer as implemented in Geneious v 11.1.5 (Biomatters, Auckland, New Zealand). For primer design, we followed the recommended criteria specified in the Fluidigm Access Array system protocol (Fluidigm, San Francisco, CA, USA), i.e. annealing temperature was set to 60°C (+/-2°C) for all primers. We targeted amplicons that were 300 to 450 bp long in order to obtain overlapping read pairs through 251bp paired-end sequencing. In order to complete this set of primers we then curated the set of 1478 single copy orthologous genes listed in Misof *et al*. (2014). Briefly, we first selected nucleotide alignments in which at least two aphid species were present (among the three present in the insect tree, i.e. *Acyrthosiphon pisum*, *Essigella californica* and *Aphis gossypii*). Among those, we then selected alignments that were longer than 800 bp, in order to work on fragments that were long enough to define conserved primers across aphids. For each of these alignments, the *A. pisum* gene sequence was retrieved and then blasted against aphid reference genomes cited above. Twenty-five genes that had blast hits against all available reference genomes were picked for primer designs, following the same procedure as explained above.

Altogether, a total of 60 pairs of primers were designed. We added Illumina linkers in their 5’ position (forward primer: 5’-TCGTCGGCAGCGTCAGATGTGTATAAGAGACAG-3’; reverse primer: 5’-GTCTCGTGGGCTCGGAGATGTGTATAAGAGACAG-3’) and tested amplification success on eight aphid DNA extracts. We chose samples from species from seven subfamilies to ensure amplification success across the aphid tree. These tests were conducted in 10 μL reaction volume using 5 µL of 2× QIAGEN Multiplex Kit Master Mix (Qiagen), 0.3 µM of each primer, and 2 µL of DNA extract. The PCR program consisted in an initial denaturation step at 95°C for 15 min, followed by 40 cycles of denaturation at 94°C for 40 s, annealing at 55°C for 45 s, and extension at 72°C for 60 s, followed by a final extension step at 72°C for 10 min. Forty-seven primer pairs that amplified all seven DNA extracts were validated. A complete list of the chosen primers as well as gene names are found in Table S3.

##### Microfluidic amplification and sequencing

We amplified these 47 markers on 96 aphid species trying to maximize overlap with species represented in the mitogenome data. We used the DNA extracts used for mitogenome amplification to which we added representatives of aphid species from additional subfamilies, tribes (Table S1); DNA extraction protocols were conducted as for aphid material produced for long-range PCRs

Microfluidic PCR were performed in an Access Array System (Fluidigm) using 2 fully loaded 48×48 Access Array Integrated Fluidic Circuits (Fluidigm): this allows for 48 samples to be simultaneously amplified across 48 distinct primer pairs. We amplified on each array the same 47 genes on 48 samples, leaving a blank template with water and no primer pair as a negative control on one of the arrays. We followed the manufacturer’s protocols except for the PCR conditions for which we used the QIAGEN Multiplex Kit Master Mix (Qiagen) and the same PCR program as for the primer tests (hybridation 30°s and 50°C) but preceded by an incubation at 50°C for 2 min and 70°C for 20 min. The PCR products were then harvested from each array for each sample (each aphid species) in 96 pools of 10 to 12μL. Eleven pools were verified on a BioAnalyzer High Sensitivity DNA Assay (Agilent Technologies, Santa Clara, California, USA). The 96 samples were PCR indexed using the UDI (Unique Dual Indexes) from Martin (2019) and the procedure described in Galan *et al*. (2018). All 96 samples were then pooled together in equal volume and 83 µL of this final pool was subjected to size selection for the full-length amplicons (expected size >350bp including primers, indexes and adaptors) by excision on a low-melting agarose gel (1.25%). It enabled to discard non-specific PCR products and primer dimers. The PCR Clean-up Gel Extraction kit (Macherey-Nagel) was used to purify the excised PCR products. It was then analyzed in a Bioanalyzer High Sensitivity Assay (Agilent Technologies, Santa Clara, California, USA) and standardized at 4nM after a KAPA qPCR quantification (KK4835; Kapa Biosystems, Woburn, Massachusetts, USA). The pooled amplicons were then submitted for paired-end sequencing on a MiSeq (Illumina) flowcell using a 500 cycle reagent cartridge v3.

##### Bioinformatic treatment

After filtering through the Illumina’s control quality procedure, reads were first demultiplexed for each aphid sample, using the sample specific dual barcode combinations and then, for each nuclear locus, using the target specific primers. We then used a custom script from Sow *et al*. (2019) to merge paired sequences into contigs with FLASH v1.2.11 and trim primers with cutadapt v. 1.9.1 Merged reads for each sample and each locus were then imported into Geneious v11.1.5. Loci which were represented by less than 25 reads were discarded from the dataset. For each locus, each aphid samples reads were first dereplicated then assembled using Geneious assembler (using the following options: high sensitivity, saving 10 contigs and generating consensus for each contig with a 85% identity threshold). The assemblies generally produced a single contig per locus per aphid sample. On a few occasions, when several contigs were obtained, one had a much higher coverage (1000 times higher than the other) and the low coverage contigs were discarded.

Gene sequences were aligned using Mafft v7.2.2.1 (Katoh and Standley 2013). Alignments were corrected to respect translation frames. Some of the genes contained intergenic regions, those were difficult to align between samples belonging to different subfamilies. The intergenic regions were therefore deleted from the final nucleic acid dataset. Finally, protein-coding genes were translated into amino acid sequences (AA). We concatenated all nucleic acid sequences to obtain a DNA matrix and all translated sequences to obtain an AA Matrix.

### *Buchnera* genome data

We used 61 published genome sequences, mainly from Chong *et al*. (2019) (16 genomes) and Manzano-Marin *et al* (2018, 2020, 2023) (29 genomes), for which taxon sampling overlapped with the one of aphid mitochondrial genomes. *Escherichia coli* K-12 MG 1655 and a selection of *Pantoea* and *Eriwnia* strains were used as outgroups. In addition, we sequenced twelve new *Buchnera aphidicola* genomes (Table S1). For both sequencing and assembly of this new dataset, we followed the protocols of Manzano-Marín *et al* (2023). Draft genomes were made of 1 to 9 contigs (Table S4) representing potentially incomplete *Buchnera*. Coding sequences (CDS) were searched in these draft *Buchnera* genomes using Prodigal 2.6.3 (PROkakyotic Dynamic programming Gene-Finding Algorithm, (Hyatt *et al*. 2010)).

The search for ortholog groups was then carried with OrthoFinder-2.5.4 (Emms and Kelly 2019) using as input Prodigal’s amino acid (AA) results for newly sequenced *Buchnera* and AA sequences from annotated published genomes. We then retrieved the single copy-core proteins of the selected genomes for phylogenetic reconstruction. We aligned the single-copy core protein sets, gene by gene, using MAFFT v7.2.2.1 (Katoh and Standley 2013) (L-INS-i algorithm). Divergent and ambiguously aligned blocks were removed using Gblocks v0.91b (Talavera and Castresana 2007). The resulting alignments were concatenated for phylogenetic inference.

### Phylogenetic analyses

The workflow of phylogenetic analyses on all three datasets is summarized in Supplementary Material Fig S1.

For both mitogenome and nuclear data, we analyzed the DNA and AA supermatrices resulting from the concatenation of all genes and implemented several models of substitution using partitioned and non-partitioned approaches. For non-partitioned analyses, we chose the best model using ModelFinder as implemented in IQ-TREE v2.1.3 (option MF) (Nguyen *et al*. 2014). For partitioned analyses, to choose the most appropriate partitioning schemes and best models, for each matrix, we built a pre-partition file: for DNA data we divided each protein– coding sequence into three partitions one for each codon position; for AA data each gene was delimited. We then ran ModelFinder (IQ-TREE v2.1.3 option MFP+MERGE) which both tests models for each partition and test if partitions can be merged. Once models and partitions were selected, we ran Maximum Likelihood (ML) analyses in IQ-TREE v2.1.3 with 1000 ultrafast bootstrap replicates (-bb option), using the best fitting models and partitions found based on BIC (Bayesian information criterion) model selection.

We also used a class of models called profile mixture models in which the likelihood of the data at each site in the alignments is calculated as a weighted sum across multiple classes, each class having its own substitution rate matrix (Pagel and Meade 2007). Modeling site-heterogeneity patterns has often been shown to outperform partition strategies in recovering tree topology (Wang *et al*. 2019), it can mitigate LBA (Long-Branch Attraction) artefacts (Lartillot *et al*. 2007) and recover contentious relationships such as relationships of bacterial endosymbionts with free-living bacteria (Husnik *et al*. 2011) or the branching of bacteria within Alphaproteobacteria (Fan *et al*. 2020). Such model is implemented in PhyloBayes for Bayesian approaches (Lartillot *et al*. 2009) and in IQ-TREE v2.1.3 (Nguyen *et al*. 2014) under ML optimization. Using Phylobayes 4.1C, we first ran the CAT model (Lartillot and Philippe 2004) on all AA matrices. For each dataset, we ran two independent Markov chains Monte Carlo (MCMC) analyses, starting from a random tree until the two chains converged (i.e. effective population size > 50 for all parameters and maximum discrepancy (maxdiff) of 0.3). Sampling trees and associated model parameters every cycle, we then computed a consensus tree with the posterior probability (pp) of each node. As stated above, IQ-TREE v2.1.3 provides a number of site-specific frequency models and profile mixture models for AA sequences, for instance C10 to C60 a variant of the CAT model applied in PhyloBayes. In order to select the best model in ML analyses, we ran the –MF on non-partitioned AA matrices including several of these mixture models (C10, C20, C50) with the -madd option in IQtree v2.1.3 and then ran an ML analysis using the best fitting models and 1000 ultrafast bootstrap. On the DNA matrices, for each dataset (mitochondrial and nuclear), we implemented a mixture of the models selected previously on the partitioned matrices using the –m Mix option. We then ran ML analyses in IQ-TREE v2.1.3 with 1000 ultra-fast bootstrap. Table S5 in Supplmentary Material lists the models implemented for each dataset under ML analyses and corresponding likelihoods of the best trees.

Contrary to the mitochondrial DNA, which is inherited as a single locus, the nuclear genes might show genealogical discordance, because of Incomplete Lineage Sorting (ILS) for instance. In order to analyze whether the lack of signal came from conflicting loci or an overall lack of information, we measured the fraction of loci and the fraction of sites consistent with a particular branch inferred in our analyses of the supermatrix (Minh *et al*. 2020). Using the ML tree presented on Fig. 3 as a reference tree, for each node, the gene concordance factor (gCF), *i.e.* the proportions of inferred single locus gene trees showing that node, were estimated. We used the options implemented in the IQ-TREE software package that takes into account variation in taxon coverage of each gene (Minh *et al*. 2020). We then estimated site concordance factors (sCF), a measure that estimates concordance at the level of sites. These node support values complements traditional bootstraps and pp values and help deciphering the cause of topological variation in phylogenetic datasets. These analyses were conducted on the nucleotide dataset as the individual gene sequences in AA were often too short to reconstruct a gene tree.

For *Buchnera* data analyses, given the high evolutionary rate of endosymbiont genomes (Moran *et al*. 1995; Wernegreen *et al*. 2001), only AA alignments were used for further analysis. The selection of substitution models on the datasets was carried out using ModelFinder (Kalyaanamoorthy *et al*. 2017) (option -MFP to test classical protein evolution models in IQ-TREE v2.1.3) using the Bayesian Information Criterion (BIC). We further performed ML searches using the selected model and an additional mixture model, C20. Bayesian inferences were conducted with the CAT model (Lartillot and Philippe 2004), as implemented in Phylobayes 4.1c (Lartillot *et al*. 2009), as explained for nuclear and mitochondrial datasets.

### Looking for sources of conflicting topologies

In order to better compare the topologies obtained with different DNA sources, we selected only species for which we obtained data from all three sources. We therefore worked on a subset of 51 aphid species that were represented by mitogenome data, nuclear genes and *Buchnera* genome data; except in two cases (namely *Rhopalosiphum* spp and *Cavariella* spp) in which for the *Buchnera* data we selected species from the same genus. We first measured statistical supports for alternative topologies using the approximately unbiased test (AU test) (Shimodaira 2002). We tested whether the AA matrix of the mitogenomes rejected or not alternative trees, *i.e*. nuclear and *Buchnera* ML trees obtained with C50 model for each respective dataset. In order to do that, trees were modified to enforce subfamily relationships observed in alternative topologies, with branch lengths all set to 1. Tests were performed on the IQ-TREE v2.1.3 with 10,000 replicates.

We then concatenated aphid nuclear and mitochondrial AA matrix in a single supermatrix and ran a new ML search using a protein mixture model (C50) implemented in IQ-TREE v2.1.3. We ran a ML analysis on a *Buchnera* dataset with the same 51 species, using also C50 substitution model. Both trees were visually compared.

We further explored whether discrepancies between aphid and *Buchnera* topology could stem from LBA effects in the *Buchnera* topology. Indeed, recent studies showed that several aphid subfamilies host very small *Buchnera*. These reduced genomes were often associated with the occurrence co-obligate symbioses (Manzano-Marín *et al*. 2023), *i.e.* the association of *Buchnera* with a secondary nutritional symbiont that complements some essential functions lost by the primary symbiont. This dual symbiotic system has been hypothesized to accelerate genome erosion in *Buchnera* (Lamelas *et al*. 2011) and could lead to convergent evolution in *Buchnera* genome, generating artefacts in the phylogeny. In order to explore this potential cause of conflicting phylogenies between *Buchnera* and its hosts, we investigated evolutionary rate shifts along branches of the *Buchnera* phylogeny. As a first test of signatures of convergent molecular evolution, we asked whether branches of the *Buchnera* tree where dual-symbiotic had evolved, underwent different rates of genome-wide molecular evolution compared to “mono-symbiotic branches”. We worked on a whole dataset limited to species where the absence/presence of dual-symbiotic systems had been investigated (Manzano-Mar^í^n *et al*. 2020; Manzano-Marín *et al*. 2018; Meseguer *et al*. 2017; Manzano-Marín *et al*. 2023; Monnin *et al*. 2020; Renoz *et al*. 2022), i.e. a subset of 60 *Buchnera* genomes. We estimated, the genome-wide ratio of non-synonymous /synonymous substitution (i.e., dN/dS) across the phylogeny reduced to this dataset, thos were inferred using the free-ratios model of PAML v.4.9e (Yang 2007).

We then used the relative rate test implemented in the R package RERconverge (Kowalczyk *et al*. 2019) (https://github.com/nclark-lab/RERconverge) to look for convergent changes in evolutionary rates in *Buchnera* genes in dual symbiotic systems. Using a species phylogenetic tree, RER converge calculates relative branch lengths for a specific gene by normalizing branch length to the distribution of branch lengths across all genes. Genes for which these standardized rates are consistently either higher or lower in foreground branches (i.e., defined as branches leading to dual symbiotic lineages) compared to background branches (i.e. all remaining branches) are identified as having experienced accelerated or decelerated molecular evolution in dual symbiotic systems. The analysis was conducted on the same set of 146 genes used for phylogenetic reconstruction. We tested for significant differences in rates within the thirteen di-symbotic lineages relative to the rest of the phylogeny. The reference phylogeny used for this analysis was the ML topology.

## Results

### Mitogenome phylogeny

Altogether, we assembled in this study 55 new complete and 33 partial mitochondrial genomes. The long-range PCR procedures generally produced one to three contigs that overlapped but did not fully recover the entire mitogenomes (Table S2). This stemmed from failures to amplify some long-range fragments. The fragment covering cox1 to cox2 was generally easily retrieved while the two other targeted fragments were amplified with more difficulties. Looking at the dataset including all complete mitogenomes, we observed very few genome rearrangement, with only a shift in the position of gene cluster *trnF-nad5-trnH-nad4-nad4l* from being flanked on its 3’-end by *trnP-nad6-cob-trnS2* to being flanked by the same gene cluster on its 5’-end as observed by Zhang *et al*. (2021) on one species of Lachninae. This shift is shared by the thirteen Lachninae and the two Thelaxinae samples for which we obtained complete mitochondrial genomes. After concatenation of the 13 coding genes of the mitogenomes, we obtained a DNA supermatrix of 11407 bp. The total AA matrix after concatenation of the 13 translated genes, was 3782 AA long.

Results of ML analyses of the concatenated AA matrix are presented on Fig. 2 with support values obtained across alternative ML analyses and PhyloBayes analyses (alternative AA topologies and topologies obtained with nucleotidic sequences are available in Supplementary Materials: Archive1). In all analyses, most subfamilies appeared monophyletic, whether AA and nuclear acid were analyzed and across substitution models. The exceptions are Eriosomatinae samples that were either scattered into three to two clades and the Calaphidinae specimens that systematically included Phyllaphidinae representatives. Depending on substitution models used for the AA matrix, Thelaxinae species either clustered within Hormaphidinae or alternatively were found as a sister group to all Hormaphidinae (PhyloBayes and ML analyses under the C50 model). Tribes represented in our dataset appeared monophyletic in all analyses except for Chaitophorini and Macrosiphini. In the latter, *Cavariella* spp. and *Muscaphis stroyani* (Smith, 1980) appeared as sister groups to all Aphidinae and were not included within Macrosiphini. Many subfamily relationships were characterized by short branches with low support values, they were also highly sensitive to substitution model specification (i.e. not retrieved in all analyses). The only subfamily relationships that were strongly sustained and consistent across analyses were: 1) the position of Neophyllaphidinae as a sister lineage to all other aphids, 2) the basal position of Lachninae; 3) the clustering of Chaitophorinae with Drepanosiphinae; 4) the clustering of Calaphidinae and Phyllaphidinae; 5) the clustering of Hormaphidinae with Thelaxinae and Anoeciinae. Additionally, a group made of Calaphidinae (including Phyllaphidinae), Chaitophorinae and Drepanosiphinae, consistently clustered with Aphidinae but with relatively low support values (bootstrap <90).

**Fig 2:**
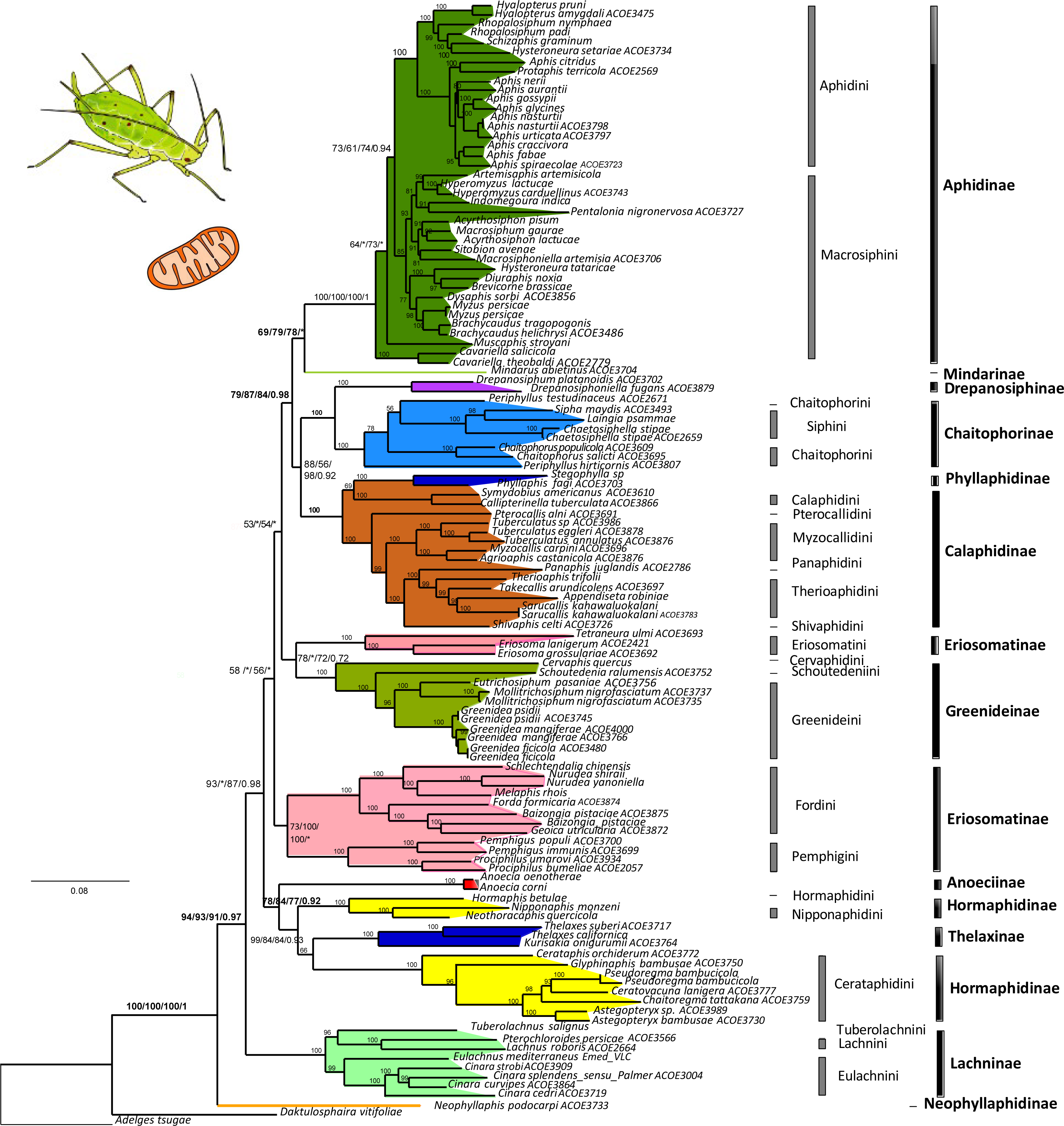
Trees obtained from the analysis of whole mitogenomes. Best tree found with ML analyses of the AA matrix under the best-fitting model (mMet+F+I+ R6) model, values at nodes indicate bootstrap values obtained with (best-fitting model)/ the C50 model/ partitioned analyses/ and posterior probabilities under the CAT model of PhyloBayes; when a “*” is indicated instead of a support values it means that the nodes was not retrieved in the corresponding analysis. Alternative support values are only given for deep nodes that were inconsistent across analyses.

### Nuclear gene phylogeny

Altogether, we obtained the sequences of 3146 amplicons across 101 aphid specimens representing 99 aphid species. Sequencing success for each locus (i.e. the number of samples for which we obtained a contig) is presented in Table S3. We failed to obtain contigs with sufficient coverage for five of the targeted loci, those were further excluded from the dataset. Once these five loci were excluded, we obtained on average a sequencing success of 70% for each amplicon (i.e. they were successfully sequenced for 70% of the species) (Table S3) and on average each species was sequenced for 27 of the 42 loci (Table S1). After concatenation of the 42 loci, we obtained a DNA supermatrix of 13400 bp, 10504 bp once intergenic regions excluded. The total AA matrix after concatenation of the 42 translated genes, was 3486 AA long. Results are presented on Fig. 3 with support values obtained across alternative ML analyses and PhyloBayes results as well as gene and site concordance factors obtained on the DNA matrix (alternative AA topologies and topologies obtained with nucleotidic sequences are available in Supplementary Material: Archive2).

**Fig 3:**
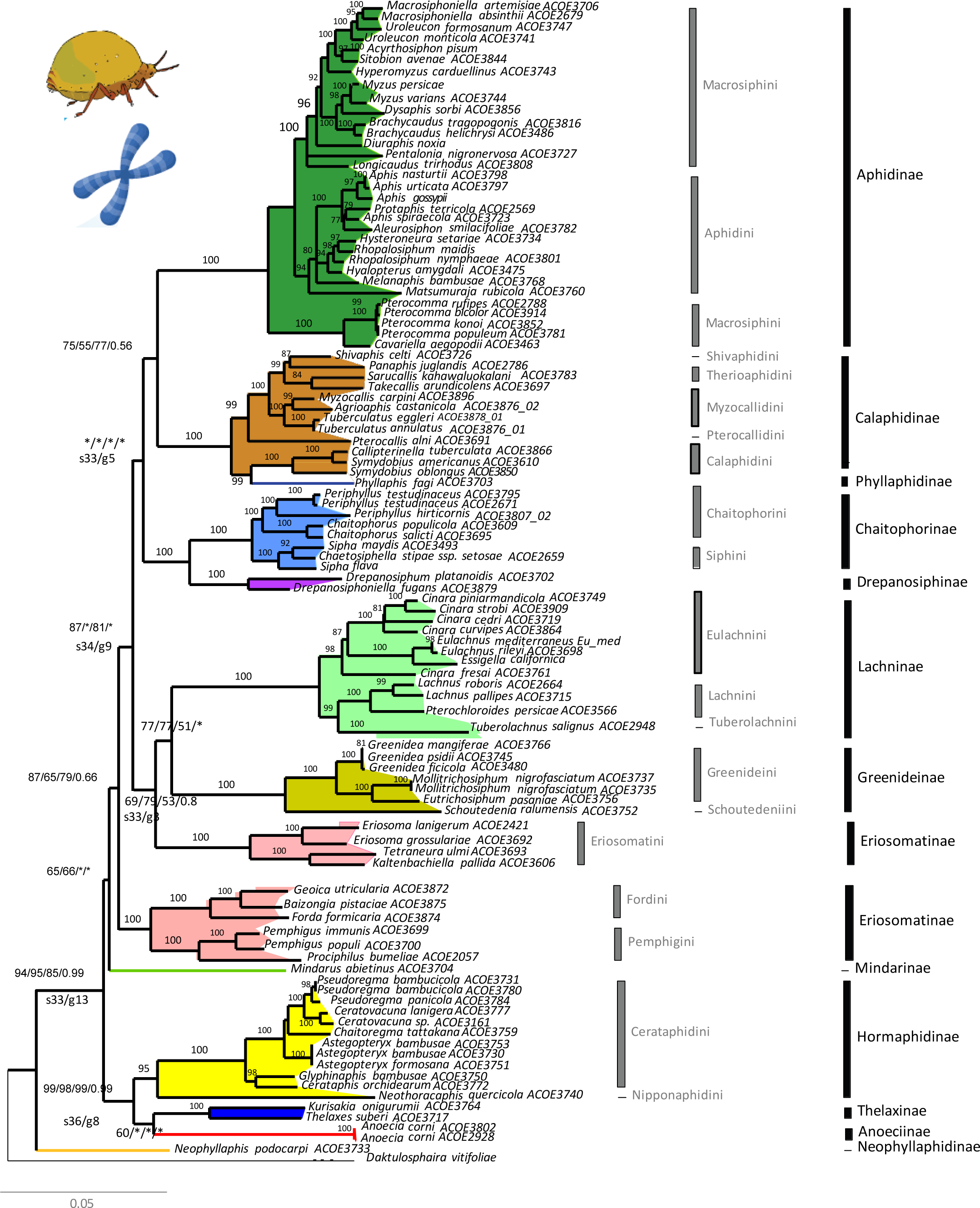
Trees obtained from the analysis of 42 nuclear loci. Best tree found with ML analyses of the AA matrix under partitioned analysis. Alternative support values are given for deep nodes: values at nodes indicate ultra-fast bootstrap values obtained under the partitioned analysis/ the non-partitioned analysis under the best-fitting model (Q.plant+F+I+R4) / the C50 model / posterior probabilities under the CAT model of PhyloBayes. When a “*” is indicated instead of a support value it means that the nodes was not retrieved in the corresponding analysis or that support was below 50.

Phylogenetic analyses confirmed the monophyly of most subfamilies with the exception of Calaphidinae that included Phyllaphidinae and Eriosomatinae which was divided into two clusters, one made of the Eriosomatini tribe, another made of Pemphigini and Fordini. Tribes were mostly monophyletic with the exception of Chaitophorini and Macrosiphini as for the mitochondrial phylogeny. Paralleling the mitochondrial phylogenetic trees: 1) Neophyllaphidinae appeared as a sister lineage of all aphids; 2) Chaitophorinae clustered with Drepanosiphinae; 3) Phyllaphidinae was positioned within Calaphidinae, 4) Hormaphidinae clustered with Thelaxinae and Anoeciinae, but their relative positions within that group were not similar across analyses (Fig. 3). Chaitophorinae, Drepanosiphinae, Calaphidinae (including Phyllaphidinae) and Aphidinae formed a clade, in ML analyses of the AA matrix under the partitioned analysis and the C50 model, but this relationship had a bootstrap support below 50 in the latter and was not retrieved in ML analysis using the Q.plant+F+I+R4 model where Calaphidinae (including Phyllaphidinae) was quite basal. However this clade was retrieved in all analyses of the DNA matrix (Supplementary Material: Archive2). The remaining subfamily relationships did not converge with the mitochondrial phylogeny and were unstable across analyses, exhibiting very short branches.

The concordance factor analyses conducted on the DNA dataset, revealed that short internal branches that generally conflicted between analyses were retrieved in very few genes (low gCF, i.e. at the most 13 of the 42 genes) and were generally supported by a sCF of 33%. The latter value is what is expected when there is no consistent signal in an alignment (Minh *et al*. 2020) (sCF are calculated for quartets, so a single site can only support one out of three topologies). The low gCf values were therefore probably due to a very weak signal in each locus and not strong gene discordance caused for instance by ILS.

### *Buchnera* phylogenetic hypothesis

We retrieved 147 highly conserved shared genes between the 73 bacterial genomes included in our analyses (i.e. 69 *Buchnera* and 4 outgroups). This low number in comparison to (Manzano-Marín *et al*. 2023) (retrieving 229 genes) likely stems from the inclusion of 12 incomplete *Buchnera* draft genomes mainly from Greenideinae (Supplementary material: Tables S4 and Archive3). After concatenation and deletion of ambiguously aligned positions; we obtained a matrix of 33820 AA. MF selected cpREV+F+I+R8 as the best fitting model. The *Buchnera* phylogeny obtained with this dataset appeared well resolved with ML and PhyloBayes analyses converging towards a very similar topology sustained by strong bootstrap and pp values (Fig. 4). As in the aphid datasets, subfamilies appeared monophyletic except for Eriosomatinae that was scattered in three or two clusters depending on analyses (i.e. ML vs Bayesian analyses, Supplementary Material Archive4) and Calaphidinae that systematically included Phyllaphidinae. As in trees based on aphid data, Chaitophorinae clustered with Drepanosiphinae while Hormaphidinae always clustered with Thelaxinae and Anoeciinae. Other relationships between subfamilies were in strong disagreements with inferences from aphid genome partitions, but they were generally robust and consistent across analyses. They generally positioned the Aphidinae subfamily (as well as Pemphigini in ML searches, Supplementary Material) as a sister group to all other aphids. The clustering of Chaitophorinae, Drepanosiphinae, with Greeneidinae and Lachninae was consistent across analyses but the relative position of Calaphidinae (including Phyllaphidinae) as well as the position of Pemphigini varied (Fig. 4).

**Fig 4:**
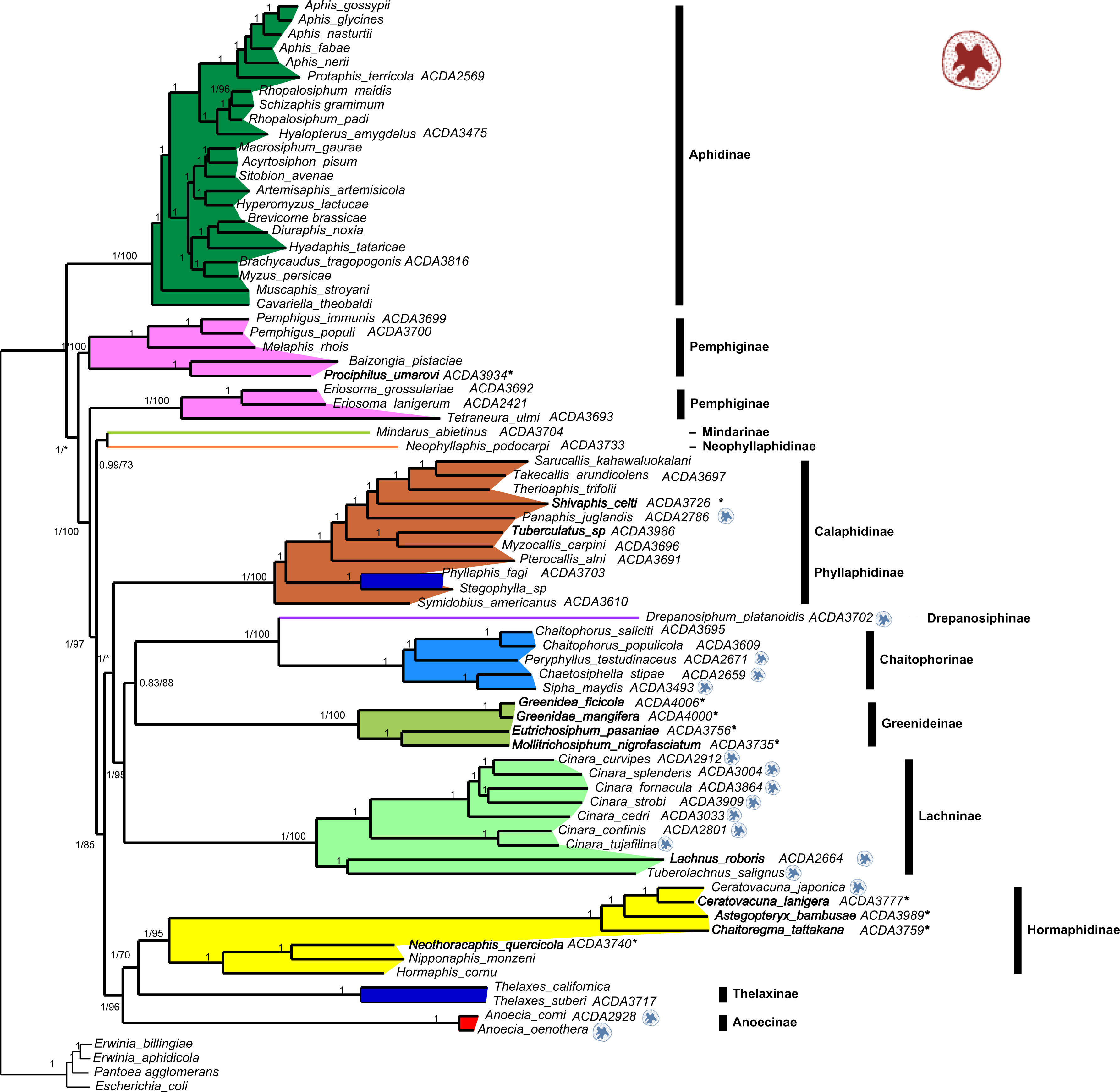
*Buchnera* tree obtained from the analysis of 146 orthologous shared single copy genes. The topology corresponds to the consensus tree found under Bayesian analyses with the CAT model of PhyloBayes: values at nodes indicate posterior probabilities in Bayesian analyses / ultrafast bootstrap values under ML analysis with the C20 model/ ultrafast bootstrap values under under ML analysis the cpREV+F+I+R8 model. * at nodes indicate bifurcations that were not observed in the ML analysis. 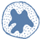 following species names indicate species for which dual-obligate symbiosis has been demonstrated, **\*** following species names indicate species for which the symbiotic status (mono-symbiotic *Buchnera* or disymbiotic system) is unknown. Names in Bold are species for which new *Buchnera* data has been acquired for this study

### Analyzing congruence and conflicts within and between datasets

When applying a congruence test (AU test in IQtree) to the mitogenome dataset, we found that the nuclear tree was not rejected (*P* = 0,247), while the *Buchnera* topology was rejected (*P* = 0.002). We then analysed a concatenated aphid data matrix (mitochondrial and nuclear) restricted to a subset of 51 aphid species that were represented by mitogenome data, nuclear genes and *Buchnera* genes.

The tree obtained from the concatenation of aphid mitochondrial and nuclear genes was very similar to the tree obtained with nuclear data only, but retrieved the clustering of Aphidinae with Calaphidinae/Phyllaphidinae and Chaitophorinae/Drepanosiphinae with good support. When compared to the *Buchnera* tree obtained on the same subset of taxa, we observed a perfect cospeciation pattern between *Buchnera* and its aphid hosts within subfamilies (or tribes in the case of Pemphiginae) (Fig. 5). On the other hand, relationships between subfamilies in the aphid and *Buchnera* tree are in disagreement except for the clustering of Thelaxinae, Hormaphidinae and Anoeciinae, the clustering of Calaphidinae and Phyllaphidinae, and the clustering of Chaitophorinae and Drepanosiphinae

**Fig 5:**
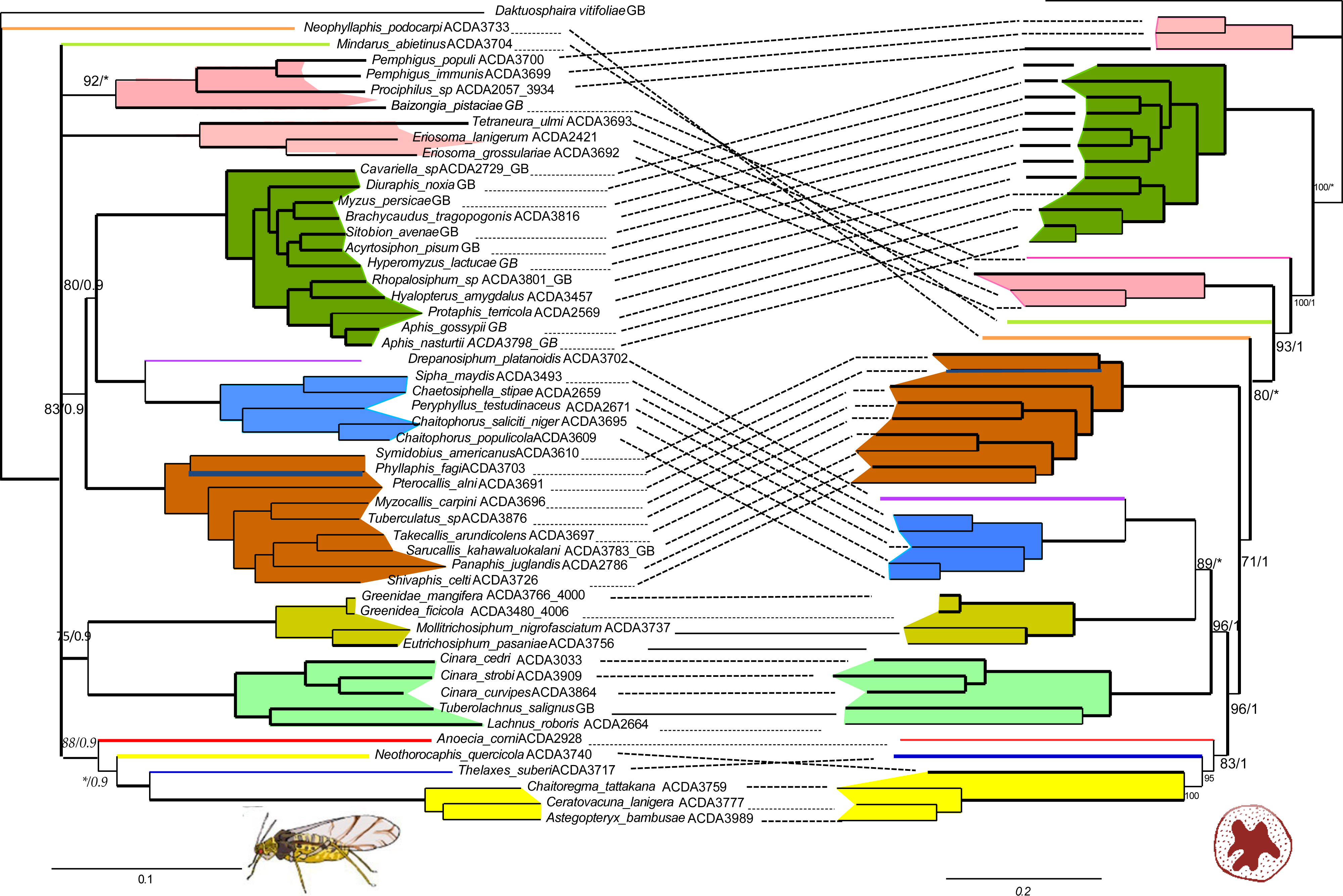
Coplot of the combined analysis of mitochondrial and nuclear data vs *Buchnera* ML tree under C50 model for species that have data from all three sources, ultrafast bootstrap values are given for subfamily relationships and pp values obtained in PhyloBayes analyses of the same dataset. Branches with support below 50 have been collapsed.

Analyses of molecular evolution patterns in *Buchnera* revealed no significant differences in overall dN/dS ratio (Supplementary Material Fig S2) between branches leading to dual-symbiotic lineages and branches leading to monosymbiotic lineages (Wilcoxon rank test, W = 429 *P* = 0.3045). Out of 138 single-copy orthologs included in the RERconverge analysis, we identified eight genes that were potentially under convergent acceleration or deceleration in dual symbiotic systems (Supplementary Material, Table S6) Our aim here was to test whether such genes could bias the results of phylogenetic analyses, we thus deleted them from the global matrix and reran an ML analysis under the C50 model. We obtained a phylogenetic tree that was identical to the tree including these genes, suggesting that they do not generate LBA in the *Buchnera* tree (Supplementary Material: Archive4).

## Discussion

Resolving the phylogeny of Aphididae is the scaffolding for a broader understanding of their evolutionary history. Previous attempts to construct a robust phylogenetic hypothesis for this family have yielded only partially satisfactory results. Mitogenomes have resolved many recalcitrant relationships across insects (e.g. Cameron *et al*. 2007; Łukasik *et al*. 2018; Chang *et al*. 2020) and relationships between lineages separated since several hundred MY (Uribe *et al*. 2019). A previous study had suggested that it could be the case for Aphididae (Chen *et al*. 2017). Nevertheless, our analyses based on a larger sampling, reveal that many internal branches remain unresolved and bear a limited amount of phylogenetic signal. Similarly, our set of nuclear genes provides restricted resolution at deeper evolutionary scales. Notably, the topologies derived from nuclear and mitochondrial datasets exhibit some incongruence. The phylogeny obtained with *Buchnera,* which genome has recurrently been used as a source of data for resolving evolutionary relationships in aphids (Novakova *et al*. 2013; Jousselin *et al*. 2009), further yields a discordant history. Although this latter phylogenetic hypothesis appears more robust than those derived from aphid DNA datasets, strong discordance between this hypothesis and previous views on the evolution of aphids raises questions about its accuracy and more generally whether a singular tree that best approximates the evolutionary relationships within Aphididae can be attained. We first discuss phylogenetic relationships retrieved across datasets in light of the taxonomic literature and subsequently explore sources of conflicts between datasets.

### Subfamily monophyly and subfamily relationships across datasets

Across all datasets, subfamilies as defined by Remaudière and Remaudière (1997) appear monophyletic except for Eriosomatinae, which is generally scattered into two groups (a cluster made of Pemphigini and Fordini and a cluster made of Eriosomatini) and Calaphidinae that always includes Phyllaphidinae. The non-monophyly of Eriosomatinae was found in previous phylogenetic studies of Aphididae (Novakova *et al*. 2013; Hardy *et al*. 2022) and studies dedicated to the subfamily (e.g. Qiao and Zhang, 2008, where Eriosomatinae is referred to as Pemphiginae). Aphids assigned to Eriosomatinae have long been set aside from other aphid groups: they share the particularity of having dwarf sexual morphs with degenerate mouthparts (Heie 1980). The unstable position of the tribes across reconstructions makes it difficult to infer the evolutionary trajectory of this characteristic, but phylogenetic reconstructions obtained so far suggest that it might have evolved several times. Eriosomatinae share a similar type of host-alternation (i.e. the occurrence of a generation of sexuparae) and the ability to form galls with Hormaphidinae, they therefore have often been suggested to be closely related to this subfamily (Ghosh 1981). But all analyses presented here reject this hypothesis, and imply that both host-alternation and gall making have evolved independently in Hormaphidinae and Eriosomatinae and maybe even several times within Eriosomatinae. The paraphyly of Calaphidinae, which in our study includes Phyllaphidinae, was retrieved in all our analyses and contrasts with the results obtained by Lee *et al*. (2022). This latter study, focusing on resolving the phylogenetic history of Calaphidinae placed Phyllaphidinae as a sister taxon of Calaphidinae, though the latter included representatives of Saltusaphidinae (not represented in our study) also rendering the subfamily paraphyletic. Though our phylogeny is based on more markers, it encompasses less species which might biases the relative position of Phyllaphidinae. In any case, the close relationships of Phyllaphidinae and Calaphidinae is consistent across studies. Because, Phyllaphidinae and Calaphidinae delimitations have varied throughout classifications, it is hard to investigate how these findings align with early views on the evolution of aphids. Our results nevertheless seem to mirror the taxonomic discussions that resulted in including “Phyllaphidinae” in a group called “Callaphididae” (see classifications of (Börner 1952; Lee *et al*. 2022)

The proximity of subfamilies Chaitophorinae and Drepanosiphinae is consistently found across our analyses. This agrees with previous aphid molecular phylogenies (Ortiz-Rivas and Martinez-Torres 2010; Ortiz-Rivas *et al*. 2004; Hardy *et al*. 2022), and more specific phylogenetic investigations within Chaitophorinae (Wieczorek *et al*. 2017). The validity of this clustering in relation to morphological investigations has been thoroughly discussed in (Wieczorek *et al*. 2017). The affinity of both subfamilies is supported by the similarity of their parasitoids (Mackauer 1965) and morphological data. Among their distinctive morphological traits, Chaitophorinae and Drepanosiphinae share the absence of sclerotisation of segment II of the rostrum, the absence of wax glands and similar male genitalia anatomy (Wieczorek *et al*. 2011; Wieczorek *et al*. 2017; Wojciechowski and Wieczorek 2004).

We also systematically recovered sister group relationships between Anoeciinae, Thelaxinae and Hormaphidinae. This again agrees with the early phylogenetic investigation of (Ortiz-Rivas *et al*. 2004) and recent phylogenomic studies (Owen and Miller 2022; Hardy *et al*. 2022). Those three aphid groups, were initially clustered in Borner’s classification within a group that also included Mindarinae and Phloeomyzinae. Apterae of all these subfamilies share a head capsule and pronotum fused with reduced (i.e. not compound) eyes (Börner, 1952). Thelaxinae and Hormaphidinae share dwarfish males (i.e. less than half the size of the female) with modified genitalia, no wings and developed mouthparts (Wieczorek *et al*. 2012). This latter character differentiates them from Eriosomatinae males that have degenerated mouthparts (Wieczorek *et al*. 2011). Hormaphidinae and Thelaxinae winged morphs also have the particularity of having wings that lie flat on the body, unlike the roof like position common among other aphids (Miyazaki 1987; Heie 1980). Hence, in light of their morphology and early taxonomical literature, the phylogenetic clustering of these three subfamilies is not surprising.

All other subfamily relationships are inconsistent across analyses. In particular, the relative positions of the two species-poor subfamilies Neophyllaphidinae (18 species) and Mindarinae (9 species), both represented by a single specimen each in our study are variable. They have been hypothesized to be sister groups of other aphids based on their association with gymnosperms (Heie 1981; Russell 1982) and their overall morphological similarities with aphid-like fossils from the upper Triassic (Heie 1967; Szwedo and Nel 2011). In particular, *Neophyllaphis* aphids closely resemble Cretaceous *Aniferella* fossils (Von Dohlen 2004). This hypothesis finds some support in our aphid data analyses for Neophyllaphidinae as *Neophyllaphis podocarpus* (Takahashi, 1920) appears as a sister taxon to all other aphids. This is also the case in the topology retrieved by Hardy *et al*. (2022). However, this basal position is not sustained in the *Buchnera* tree. Concerning Mindarinae, its position is more variable across trees. Finally, Lachninae were found at a basal position in the mitochondrial tree whereas they appeared related to Greenideinae in both *Buchnera* and aphid nuclear DNA trees. No a priori can be found in the taxonomic literature on the relatedness of these two subfamilies, but interestingly they show similarities in male genitalia morphology (Wieczorek *et al*. 2012).

Below the subfamily level, our analyses retrieved the monophyly of all the tribes represented except for *Macrosiphini*, as *Cavariella*, *Pterocomma* genera and *Muscaphis* are positioned as sister groups of a clade made of Aphidini and the remaining *Macrosiphini*. The non monophyly of Macrosiphini was already suggested by (Novakova *et al*. 2013; Hardy *et al*. 2022) and a study dedicated to Macrosiphini (Choi *et al*. 2018). Using a very different sampling and set of DNA markers, all three studies show that Macrosiphini are not monophyletic and that *Cavariella* and/or *Pterocomma* (depending on species sampling) are allied and form a separate clade probably with several other Aphidinae genera.

Altogether, our reconstructions validate the classification of aphids into subfamilies as proposed by Remaudière and Remaudière (1997). This alignment with the existing classification is unsurprising, given its recognition as a consensus framework. Concerning tribe delimitations, as exemplified by our results in Macrosiphini, a thorough investigation of each subfamily, with dense sampling in each tribe will be needed to reach firm conclusions. Beyond the subfamily level, our results offer limited support for higher-level phylogenetic groupings, apart from the three clusters discussed above. This overall lack of resolution parallels the historical trends within the taxonomic literature, marked by controversies surrounding higher-level classifications and aligns well with the idea that aphids have undergone a rapid radiation (von Dohlen and Moran 2000; Von Dohlen 2004) as supported by the fossil record (Heie 2004; Szwedo and Nel 2011).

### Conflicts between datasets: is *Buchnera* telling the truth?

*Buchnera* phylogeny is fully resolved but it conflicts with the aphid topology. There is one striking disagreement between aphid and *Buchnera* topologies: the relative position of Aphidinae. In the *Buchnera* phylogenetic hypothesis, this subfamily appears as a sister group to other aphid subfamilies while in aphid datasets, it appears in a more derived position (it clusters with Chaitophorinae/ Drepanosiphinae and also Calaphidinae/ Phyllaphidinae in most analyses). These results either imply that: 1) *Buchnera aphidicola* has not systematically cospeciated with its aphid host; 2) the aphid phylogenies are too poorly resolved in their deep nodes to be used as solid phylogenetic hypotheses; 3) *Buchnera* phylogeny is plagued by artefacts and does not accurately reflect aphid subfamily relationships.

The first hypothesis goes against all evidence accumulated in the literature on *Buchnera* vertical transmission (e.g. Wilkinson *et al*. 2003; Hinde 1971; Buchner 1965b; Koga *et al*. 2012) and previous phylogenetic evidence of aphid/ *Buchnera* cospeciation at different evolutionary scales (*e.g.* Clark *et al*. 2000; Jousselin *et al*. 2009; Xu *et al*. 2018). Moreover, the current study also shows that within subfamilies, *Buchnera* and aphid phylogenetic histories are perfectly congruent (Fig. 5). The lack of cospeciation between aphids and *Buchnera* in deep nodes, would imply that horizontal transfers if any, have only occurred during the early diversification of aphids or that *Buchnera* was acquired independently in different aphid lineages. This latter hypothetical scenario, would supposes that aphids had initially diversified fixing different nutritional symbionts (as observed in aphid sister group Adelgidae; von Dohlen *et al*. 2017), recurrently recruiting a *Buchnera*-like endosymbiont (as observed for *Serratia symbiotica* as a co-obligate partner of aphids, Manzano-Marín *et al*. 2023). Current aphid lineages would be the ones that kept *Buchnera*, while others became extinct. While this alternative appears more biologically realistic than horizontal transfers of *Buchnera*, it still seems unlikely considering that almost all aphids host *Buchnera* and that those are almost perfectly syntenic (Chong *et al*. 2019; Manzano-Marín *et al*. 2023). This synteny is never observed in study systems with multiple symbiont acquisitions (Dial *et al*. 2021) and is good indicator of *Buchnera* single acquisition event. Given uncertainties in host and symbiont trees, we favor a scrnerio in which lack of convergence in host and symbiont trees is caused by phylogenetic irresolution andwe have knowingly chosen not to conduct reconciliation analyses between aphids and *Buchnera* to and explore horizontal transfers and multiple acquisition scenarios.

An alternative hypothesis for the lack of parallelism of aphid and *Buchnera* phylogenies is that deep nodes in the aphid trees are unreliable while *Buchnera* gives a more solid phylogenetic hypothesis. As mentioned previously, the mitochondrial deep nodes are poorly resolved. In addition, our tree is very different from the one obtained by Chen *et al*. (2017) though we employed very similar analytical tools but on an extended taxon sampling. This suggests that phylogenetic resolution using mitogenome data is very sensitive to taxon sampling which usually indicates difficulties in estimating substitution models (Bernot *et al*. 2023). The nuclear DNA data does not yield a more stable hypothesis. On the other hand, *Buchnera* topology is consistent across all analyses and both Chong *et al*. (2019) and Manzano-Marín *et al*. (2023), conducted on a smaller sampling, yield very similar hypotheses about subfamily relationships. *Buchnera* evolves fast (Moran *et al*. 1995; Moran 1996) as other primary symbionts of arthropods (Woolfit and Bromham 2003; Degnan *et al*. 2004; Arab *et al*. 2020). If a rapid radiation has indeed given rise to aphid subfamilies, there might have been limited time for informative substitutions to be fixed between speciation events in the aphid genome while the fast evolutionary rate of *Buchnera* might have allowed their retention. Following this argument, the *Buchnera* phylogeny could represent a more solid phylogenetic hypothesis than aphid phylogenies.

However, the affinity of Aphidinae with Calaphidinae/ Phyllaphidinae and Chaitophorinae/ Drepanosiphinae is also observed in two recent phylogenomic investigations (Owen and Miller 2022; Hardy *et al*. 2022). Though it is hard to be fully confident on the robustness of this result (the support is low in Owen *et al*.’s study and it is not given in Hardy *et al*. 2022), its consistency across studies suggests that it is not caused by a sampling bias or limited nuclear marker sampling. Recent investigation on male morphs and male genitalia, also validates this clustering (Wieczorek *et al*. 2011): all five subfamilies share normal sized males without modified genitalia. Furthermore, in the taxonomic literature, Aphidinae is consistently considered as a more derived subfamily. This stems from its scarcity in the fossil record, as well as morphological characters and biological characteristics such as associations with Rosaceae and diversification on herbaceous hosts (Shaposhnikov 1981; Heie 1981). Though some of these arguments can be discussed, they cast doubt on the hypothesis that *Buchnera* “is telling the truth“, *i.e.* proposing a solid phylogenetic hypothesis for its hosts. There remains the possibility that some properties of *Buchnera* genomes alter phylogenetic reconstruction efforts. *Buchnera* genomes exhibit strong compositional biases. The use of mixture substitution models is supposed to alleviate the problem of treating such data (Lartillot and Philippe 2004; Lartillot *et al*. 2007; Rodriguez-Ezpeleta *et al*. 2007). However, we have no estimate of their absolute goodness of fit to our data and they can remain inadequate and still too simplistic compared to the evolutionary process driving endosymbiont molecular evolution. In addition, the multiple occurrences of dual-symbioses (Manzano-Marín *et al*. 2023) might further impact *Buchnera* genome evolution. Though our tests for shifts in evolutionary rates between branches of the *Buchnera* tree were not significant, our analyses are preliminary and exclude certain aphid lineages groups which symbiotic status is uncertain (Greenideinae and some Hormaphidinae) (Fig. 4). In Hormaphidinae, we have evidence that at least one species has a co-obligate symbiont (Yorimoto *et al*. 2022) and some species have completely lost *Buchnera* (Fukatsu *et al*. 1994; Vogel and Moran 2013). In Greenideinae, the occurrence of a dual symbiotic system has not been investigated, but the *Buchnera* draft genomes presented here show a highly biased composition which could indicate further *Buchnera* degeneration (Table S4). More information on the occurrence of dual-symbiotic systems across and within subfamilies is therefore needed to conduct thorough tests of evolutionary rate shifts in *Buchnera*. In any case, given *Buchnera* erratic pattern of genomic erosion, we cannot confidently affirm that it can be relied on to reconstruct robust evolutionary scenarios about aphid subfamily relationships.

### Conclusions and perspectives

Heie stated in 1980 that there were “*as many aphid classifications as taxonomists*”. Paraphrasing this, we could say that there are “*as many phylogenies as phylogeneticists*”. Can we obtain a more solid phylogeny with more data, i.e. more species with a representation of all subfamilies/ tribes and genome scale data? We can hope so. However, genome scale data can also increase the probability of observing incongruent signals. In addition data errors, miss-assembly and contaminations often plague phylogenomic analyses (Jeffroy *et al*. 2006; Simion *et al*. 2020). When no a priori hypotheses on phylogenetic relationships are expected, as in aphids, such artefacts are hard to track. Therefore we want to urge for caution in interpretation of future aphid phylogenomic hypotheses. Their validity in light of morphological characters, will need to be evaluated. Interestingly, all strongly sustained subfamily relationships in our study are supported by shared male genitaliae morphology. Therefore, as underlined by Wieczorek *et al*. (2012), male morphology might actually be a good marker of aphid relatedness. The advent of chromosome level assemblies could also bring new phylogenetic information such as chromosomal architectural changes (as in Schultz *et al*. 2023). Finally, if the aphid initial diversification is actually the result of a fast radiation (von Dohlen and Moran 2000), as for numerous other groups such as birds to cite one that is not short of phylogenomic investigation (Suh 2016; Reddy *et al*. 2017), their early diversification will be difficult to solve.

On a positive side, we consolidate several results obtained in previous studies encompassing less subfamilies and/or using less DNA data sources: the clustering of Drepanosiphinae with Chaitophorinae, the clustering of Calaphidinae with Phyllaphidinae, the clustering of Hormaphidinae, Thelaxinae and Anoeciinae, the paraphyly of Eriosomatinae and its lack of phylogenetic affinity with Hormaphidinae. Given these relationships, we confirm that two aphid biological characteristics considered as complex-host-alternation (present in Anoeciinae, Hormaphidinae, Eriosomatinae and Aphidinae) and gall forming (present in Eriosomatinae and Hormaphidinae)-have evolved independently in several aphid subfamilies. Finally, we have developed here a useful set of nuclear DNA markers that appear useful for solving relationships within aphid subfamilies. This means that some of the questions that still hover in aphid evolutionary biology -evolution of gall making, role of host plant shifts and biogeographic history in species diversification, evolution of dual-symbioses-can be addressed in a strong phylogenetic framework within some of the highly diversified aphid subfamilies.

## Supporting information

Sulementary tables

Archive1

Archive2

Archive3

Archive4

## Contribution statement

E.J.: conceptualization, formal analysis, investigation, writing--original draft, funding acquisition. A.CDA: conceptualization, investigation, writing—review and editing. ALC.: investigation, writing— review and editing. MG: conceptualization, investigation, writing—review and editing. CC : investigation, writing—review and editing. VB: investigation, writing—review and editing. AMM conceptualization, formal analysis, investigation, writing--review and editing, funding acquisition.

## Data Availability Statement

The analysed datasets underlying this article are available in Zenodo and can be accessed at https://zenodo.org/uploads/10513209. ENA or NCBI accession numbers for mitogenomes and *Buchnera* genomes are available in Table S1.

## Acknowledgements

We would like to acknowledge the talented artist/scientist Jorge Mariano Collantes Alegre for the aphid cartoons. This work was supported by the Marie-Curie AgreenSkills+ fellowship programme co-funded by the EU’s Seventh Framework Programme (FP7-609398) to A.M.M, the Marie-Skłodowska-Curie H2020 Programme (H2020-MSCA-IF-2016) to E.J., the France Génomique National Infrastructure, funded as part of the *Investissement d’Avenir* program managed by the *Agence Nationale pour la Recherche* (ANR-10-INBS-0009) to E.J., C.C., and V.B and INRAe department SPE project “Impact Phyto” (2016-2019) to EJ and Gael Kergoat. We are grateful to the genotoul bioinformatics platform Toulouse Midi-Pyrenees (Bioinfo Genotoul) and the CBGP-HPC computational platform for providing help and/or computing and/or storage resources. Data used in this work were (partly) produced through the GenSeq technical facilities of the «Institut des Sciences de l’Evolution de Montpellier » with the support of LabEx CeMEB, an ANR “Investissements d’avenir” program (ANR-10-LABX-04-01). The funders had no role in study design, data collection and analysis, decision to publish, or preparation of the manuscript.

