## Supplementary figures and images for "Discordance between phylogenomic datasets in aphids: who is telling the truth?"

### AA_concat_QplantFluidigm.contree.pdf

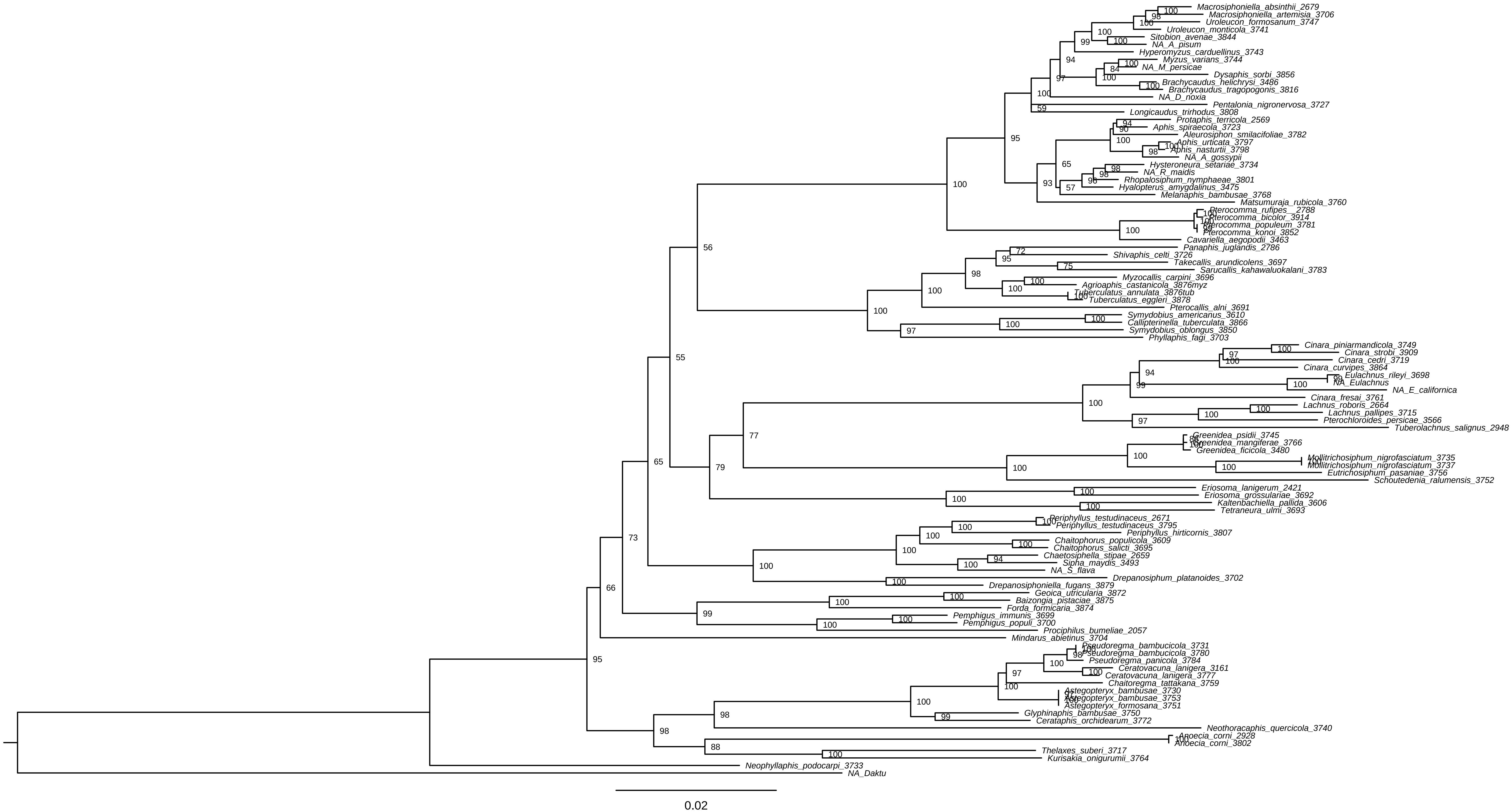

### AAIQtreeC50FluidNew.pdf

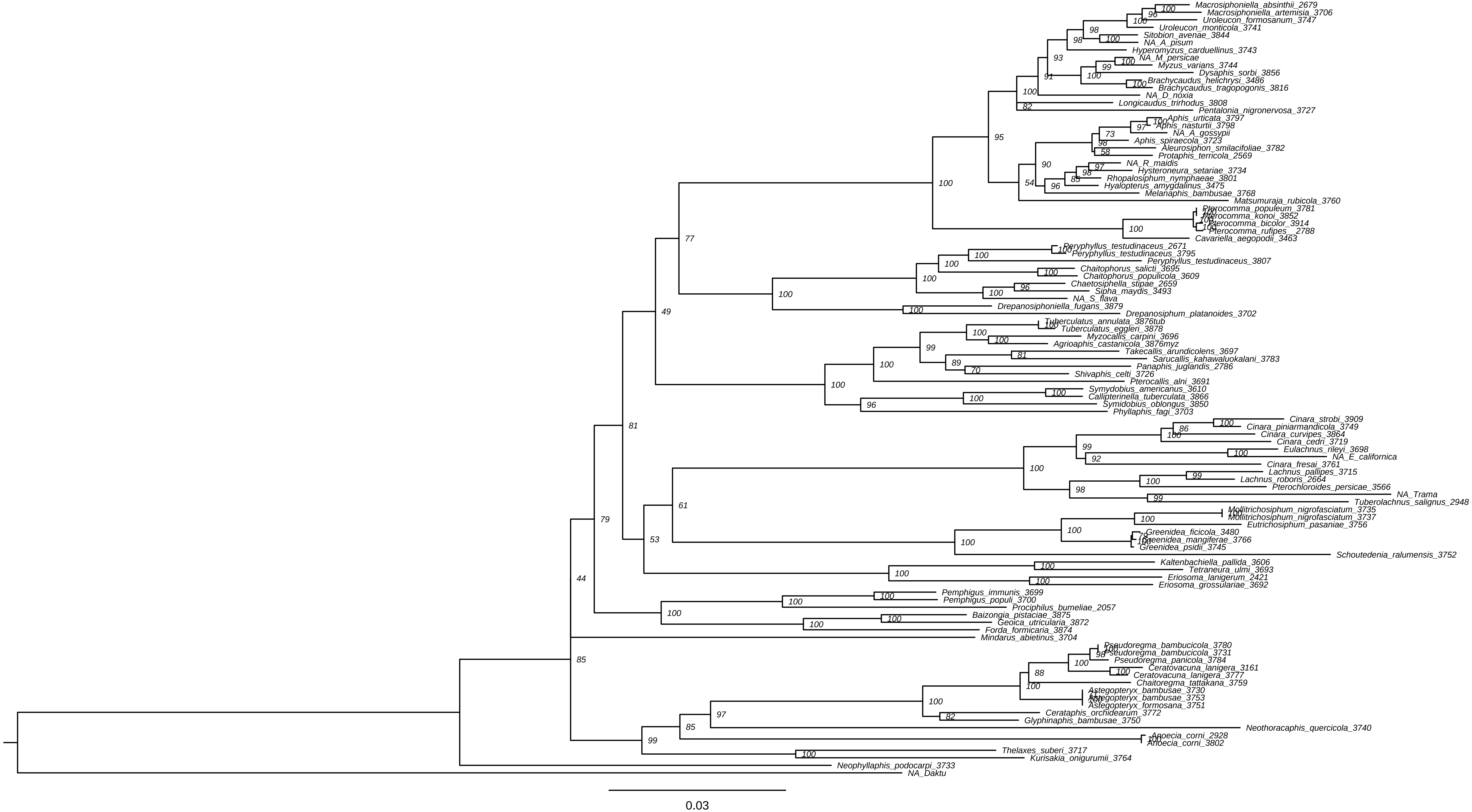

### AAIQtreePartitionFluidNew.pdf

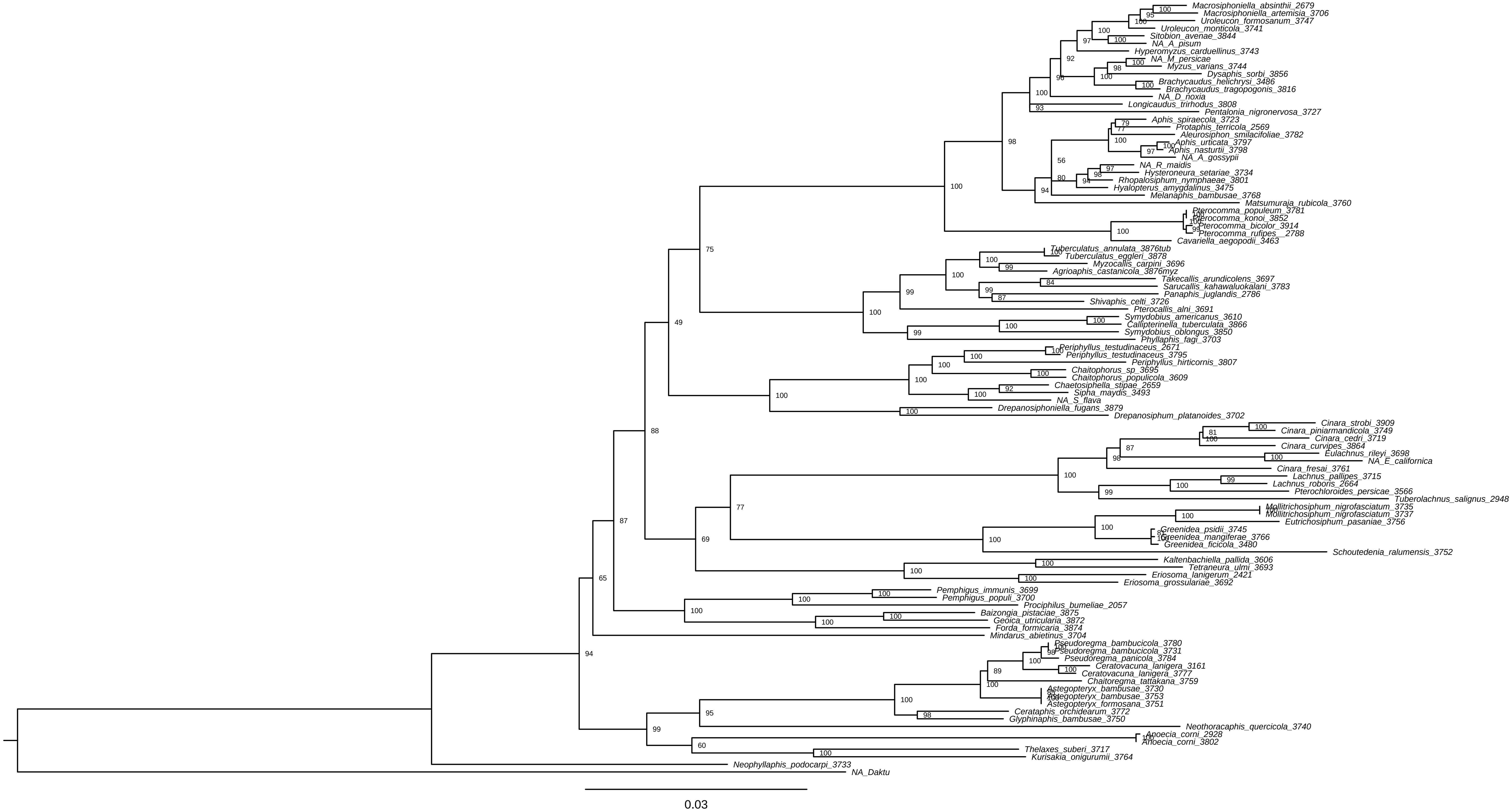

### BuchneraMLTreeC20NoRERGene.pdf

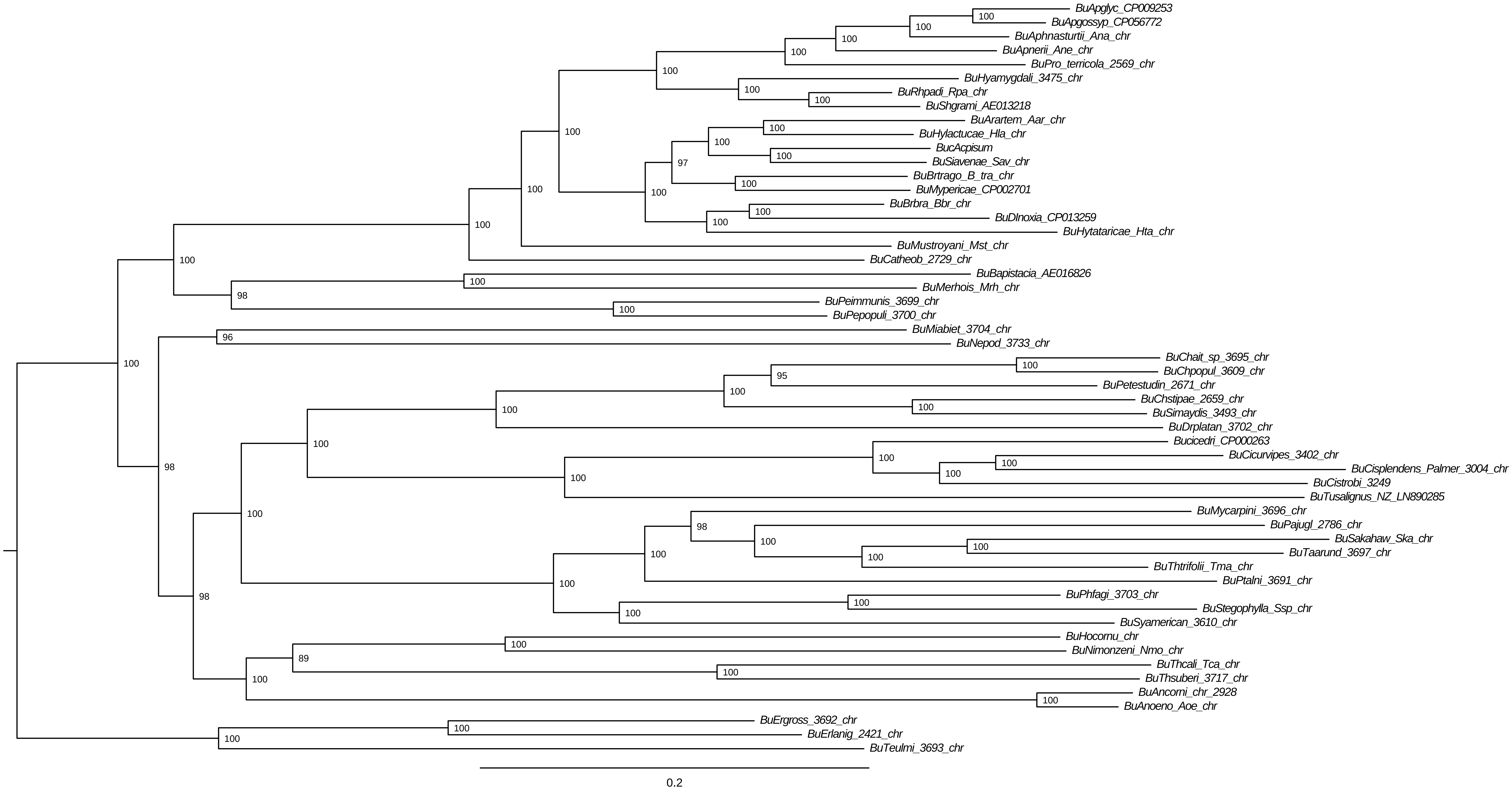

### C50_AA_MitochConTree.pdf

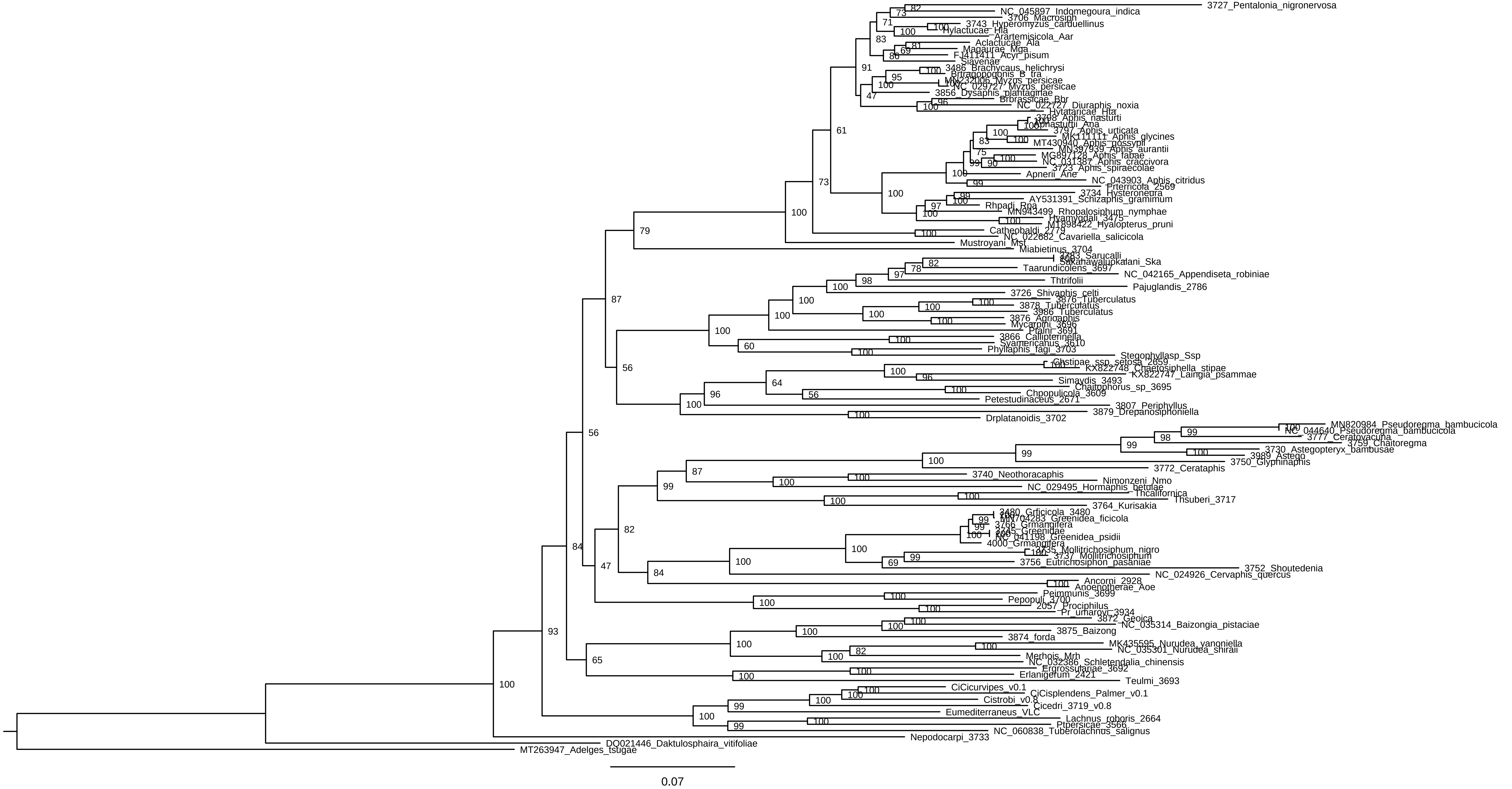

### FigureS1

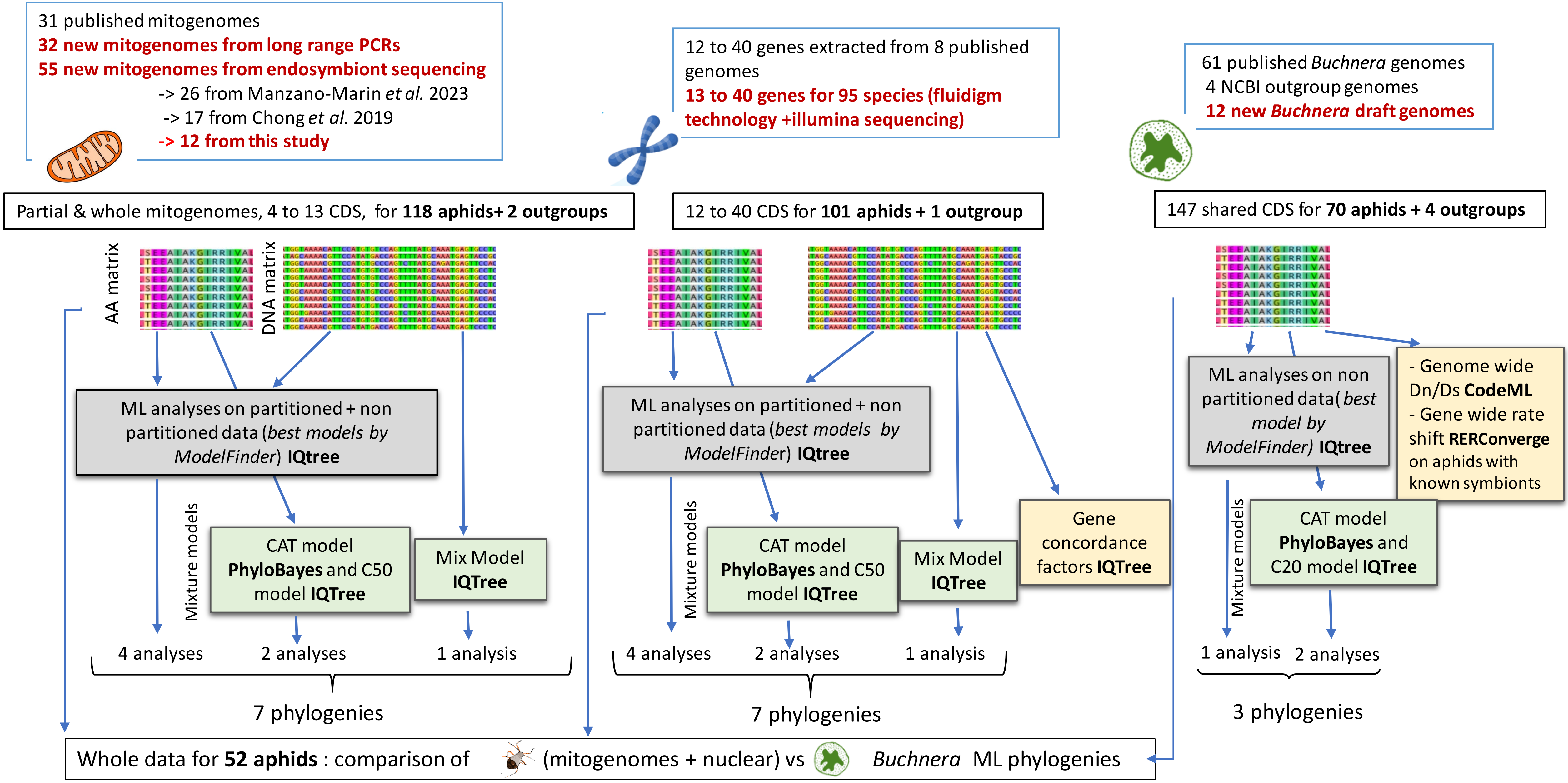

### FigureS2

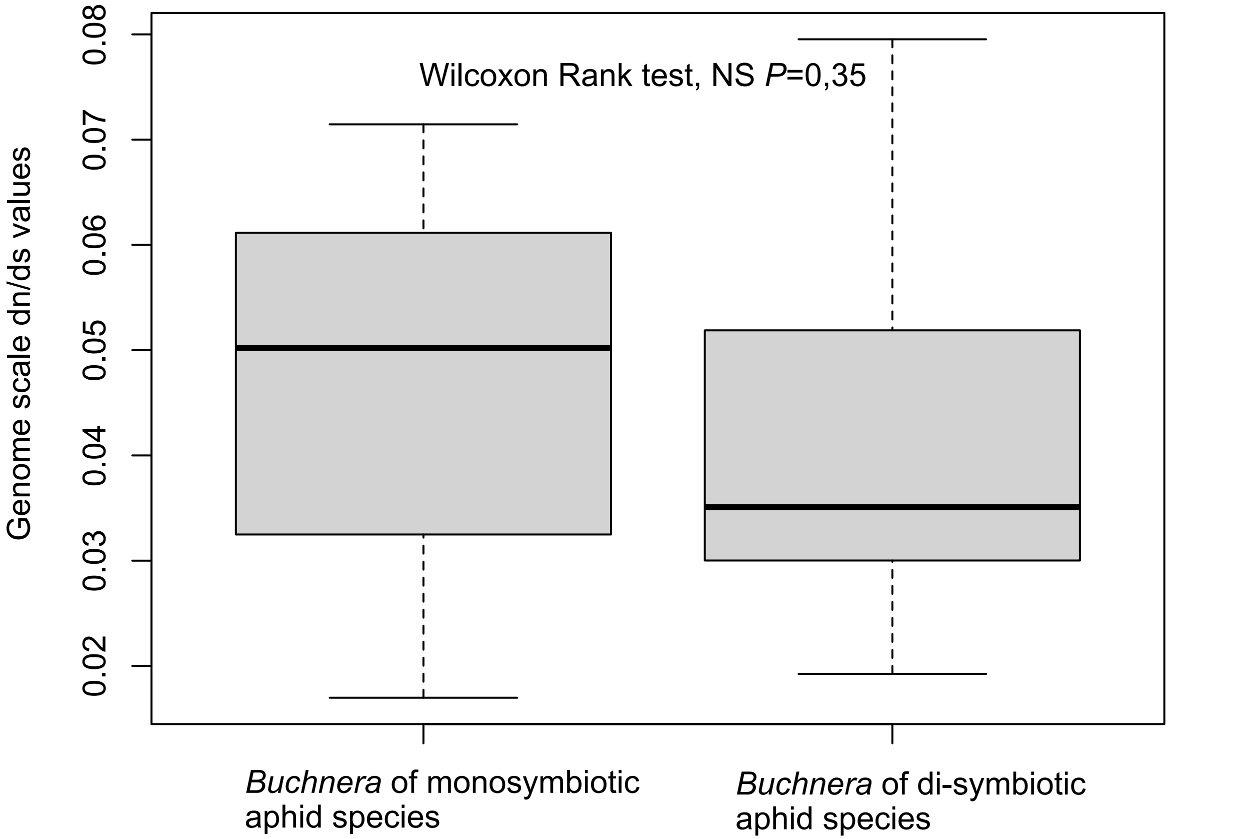

### Met_AA_Mitoch_contree.pdf

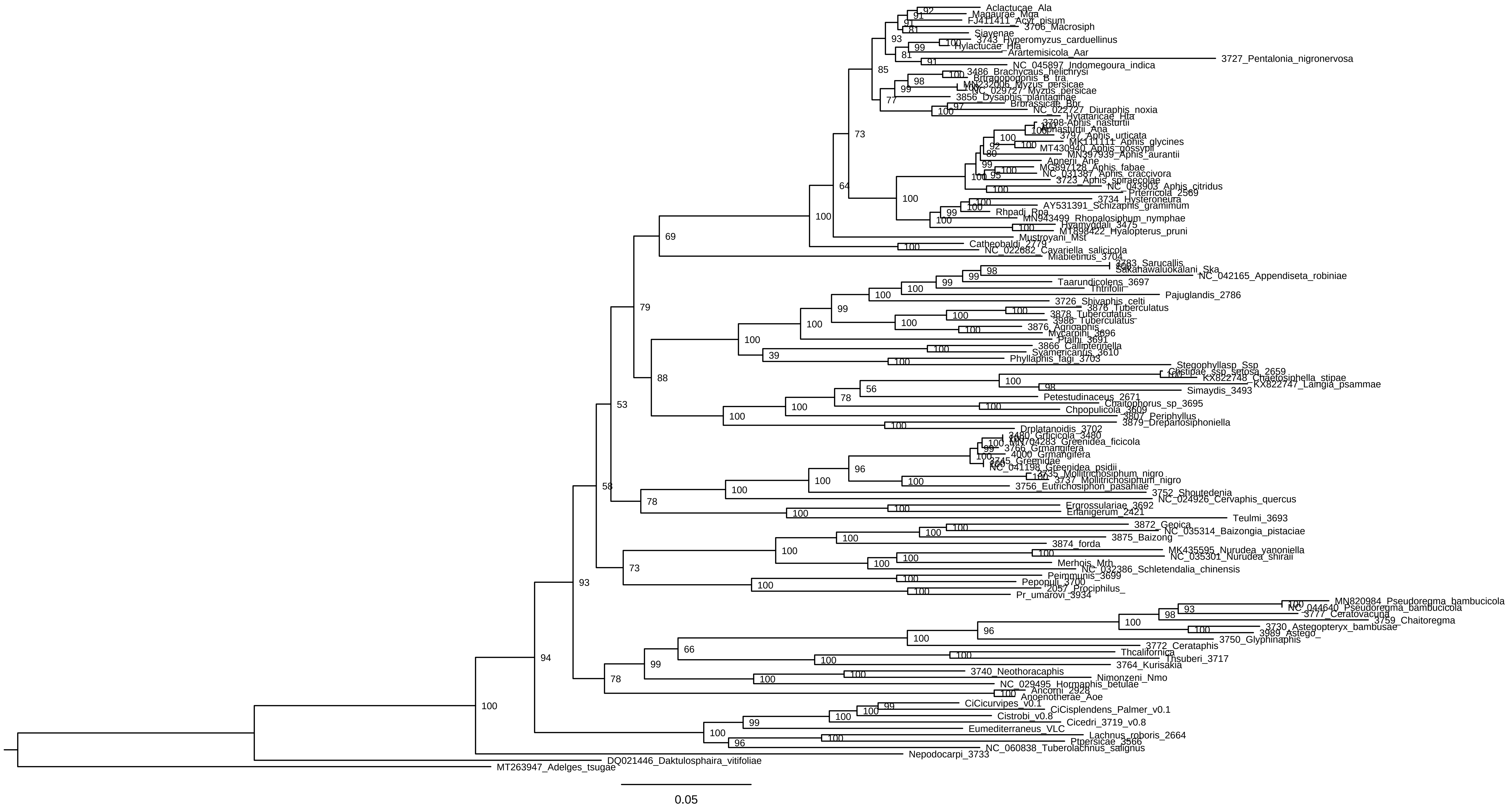

### ML_cpREV+F+I+R8_Buchnera.pdf

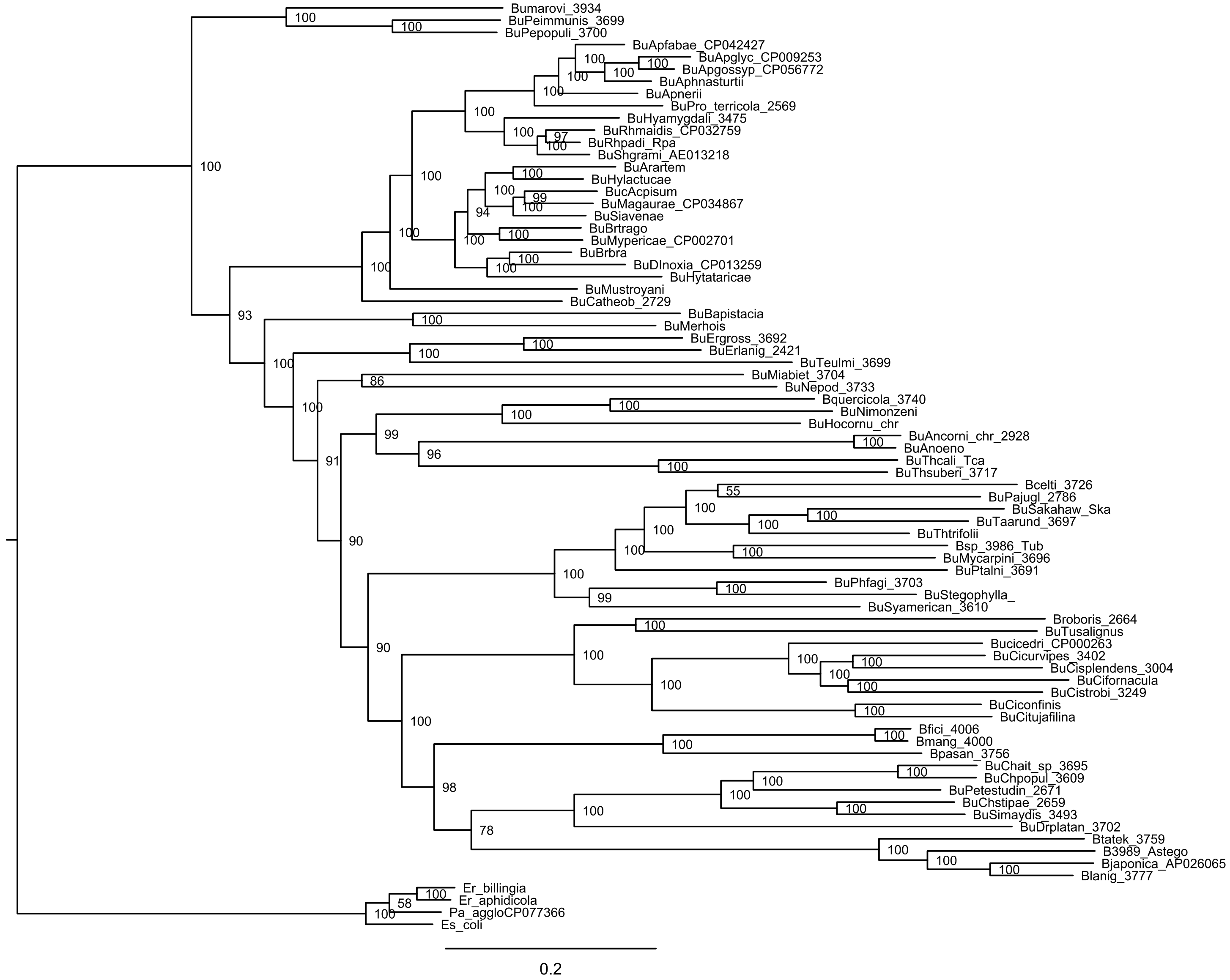

### MLC50BuchneraTree.pdf

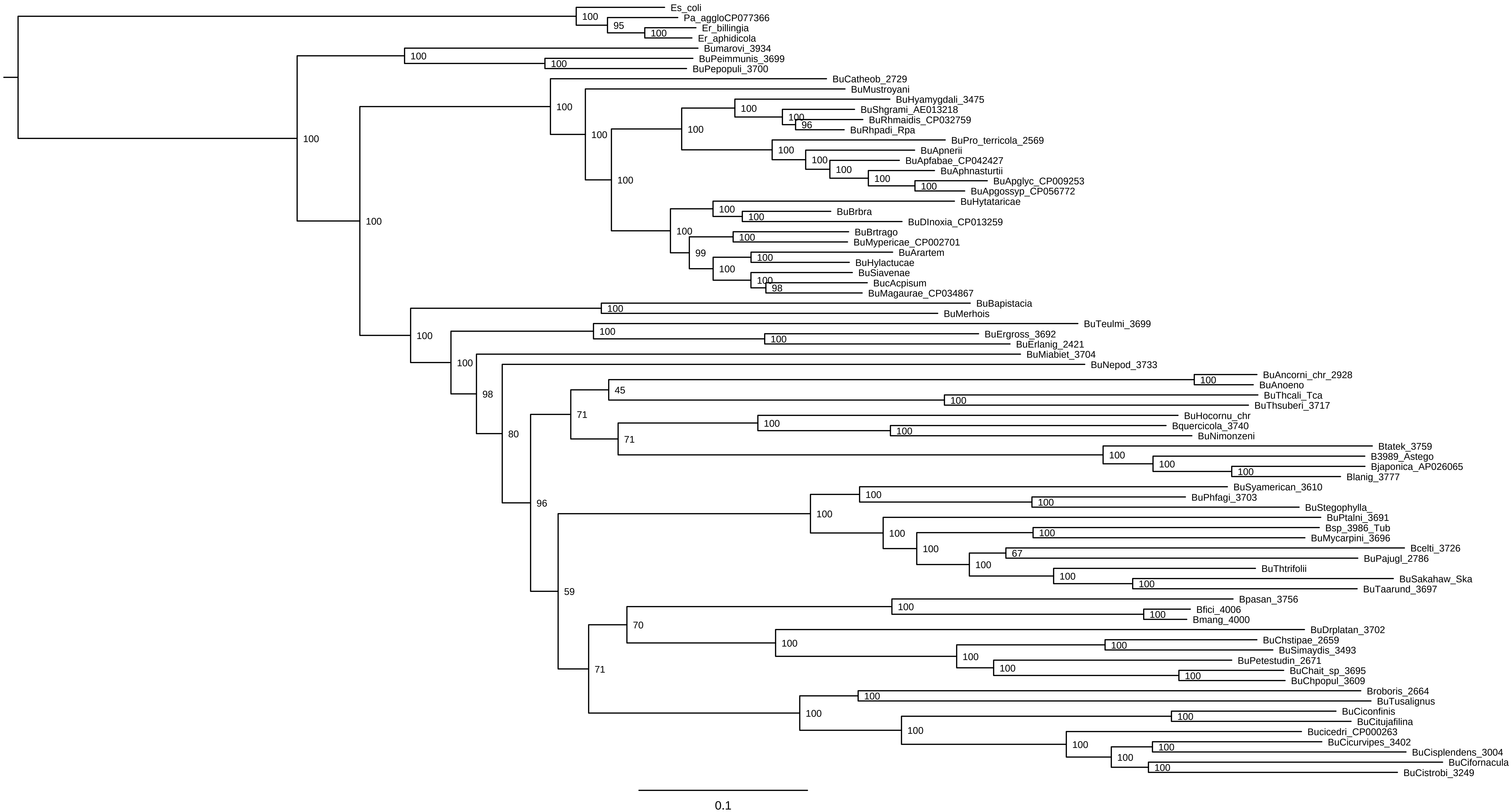

### Nuc_short_2024_GTR_fluidigm.pdf

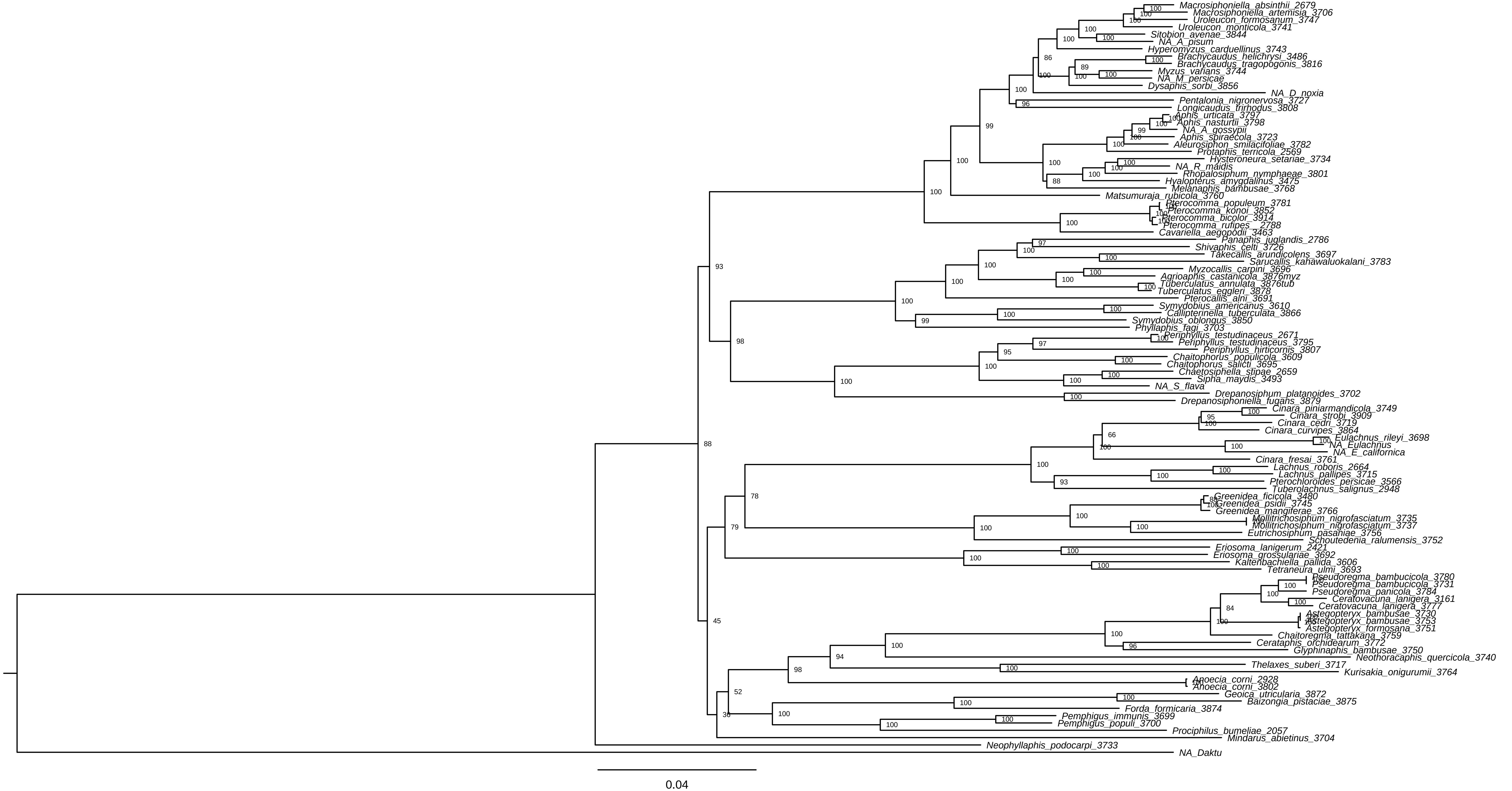

### Nuc_short_2024_Mixture_fluidigm.pdf

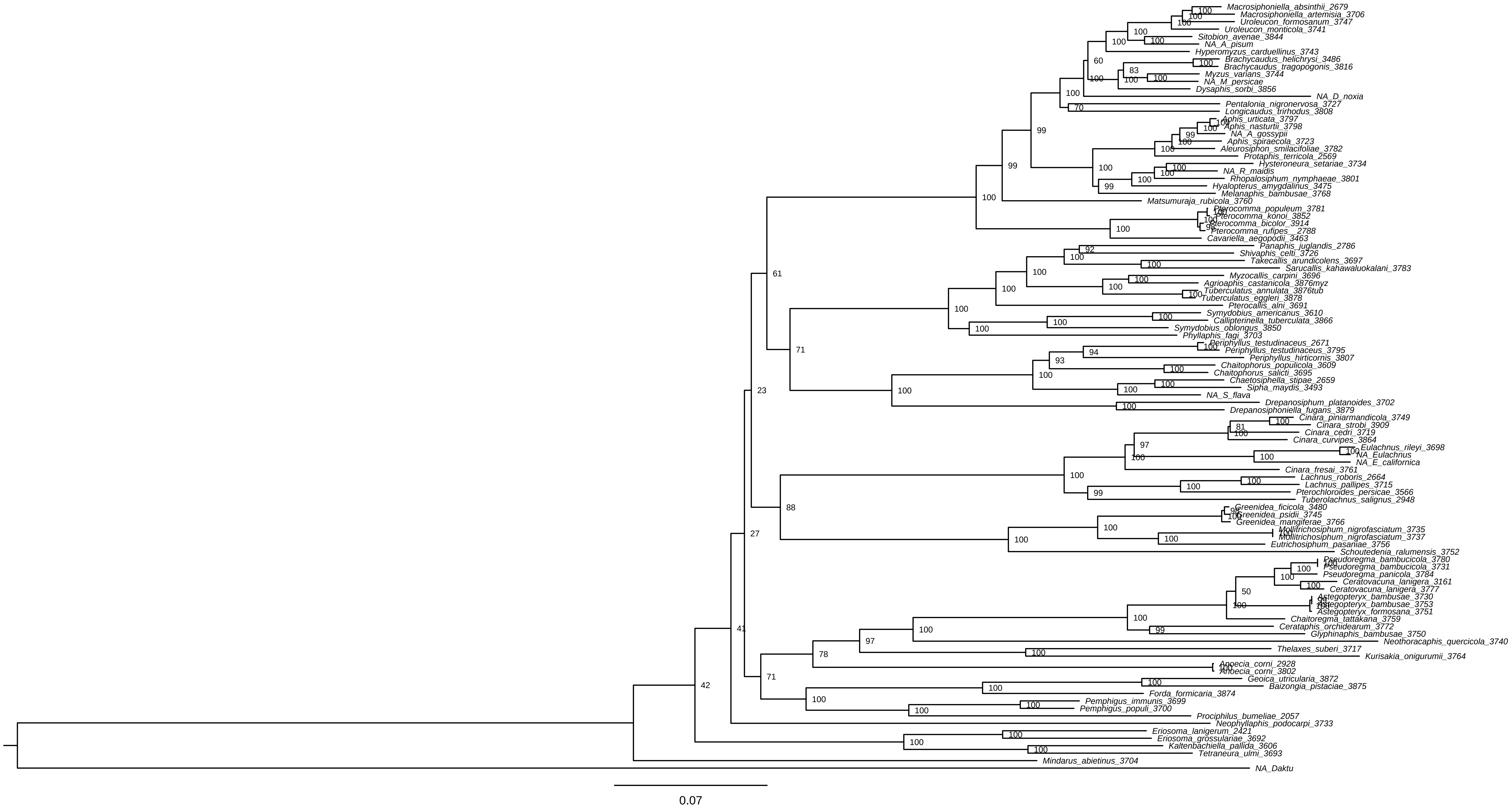

### Nuc_ShortPartition.pdf

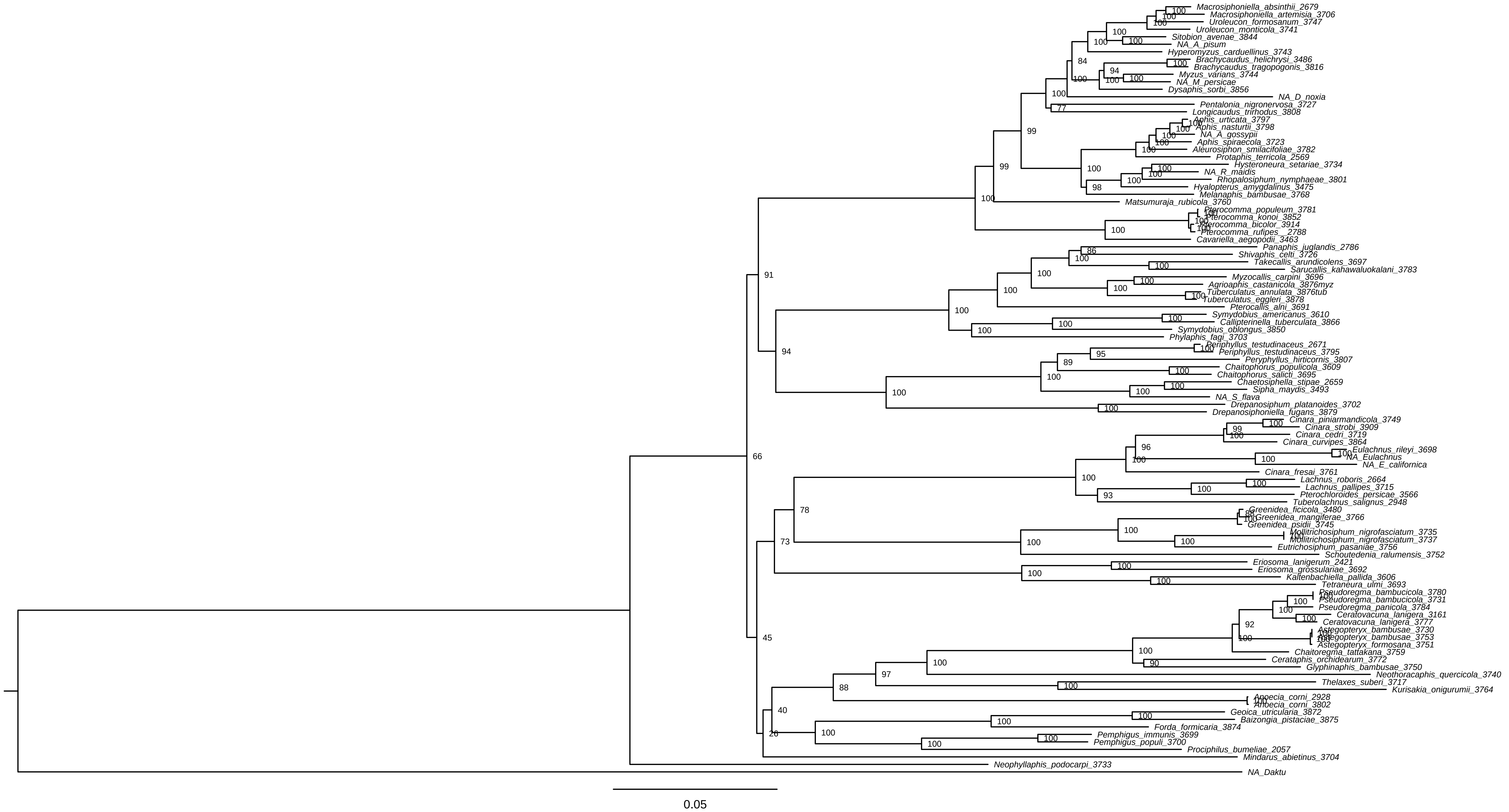

### NucMitoch_C50_contree.pdf

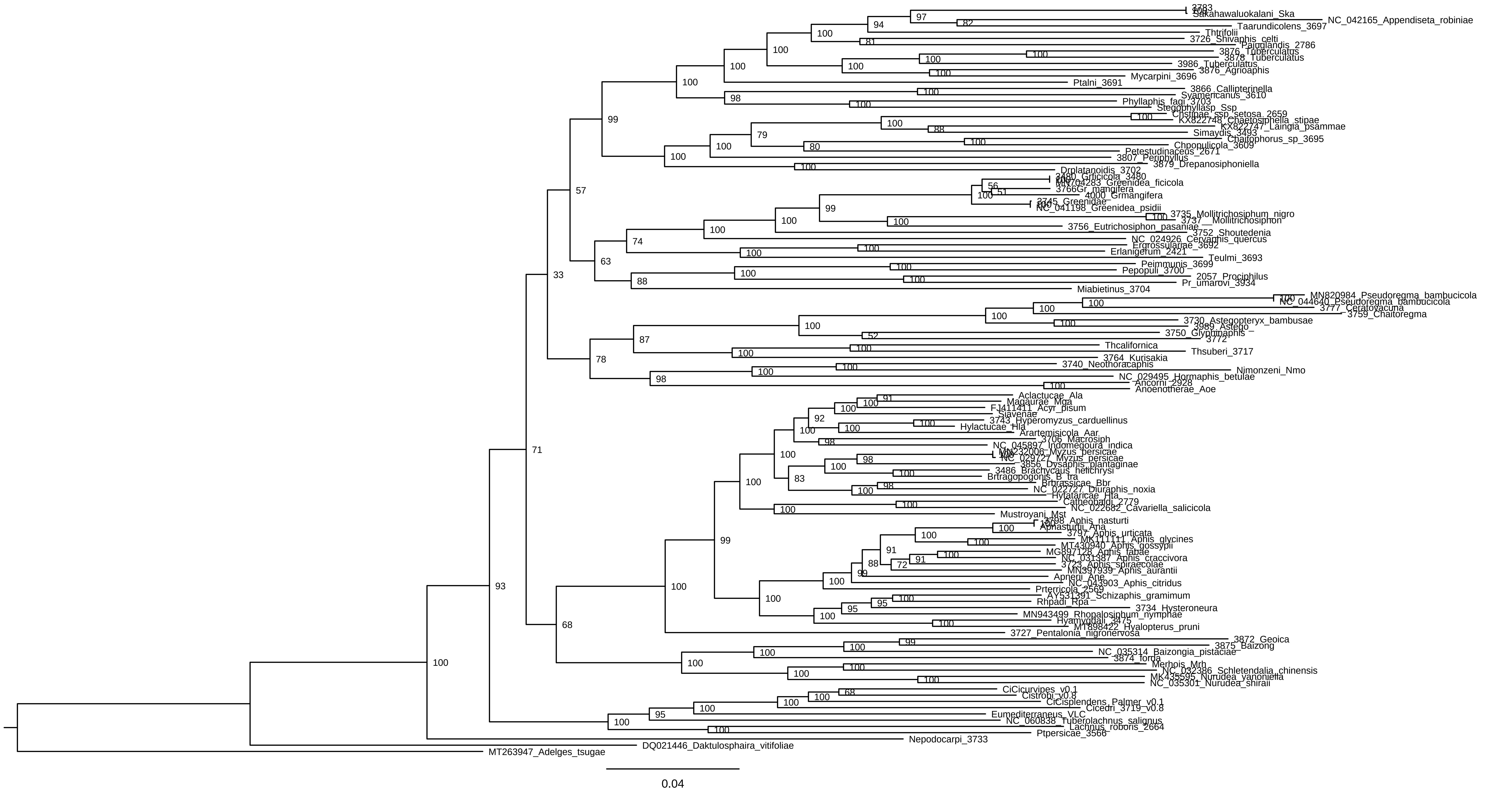

### NucMitoch_GTR_conTree.pdf

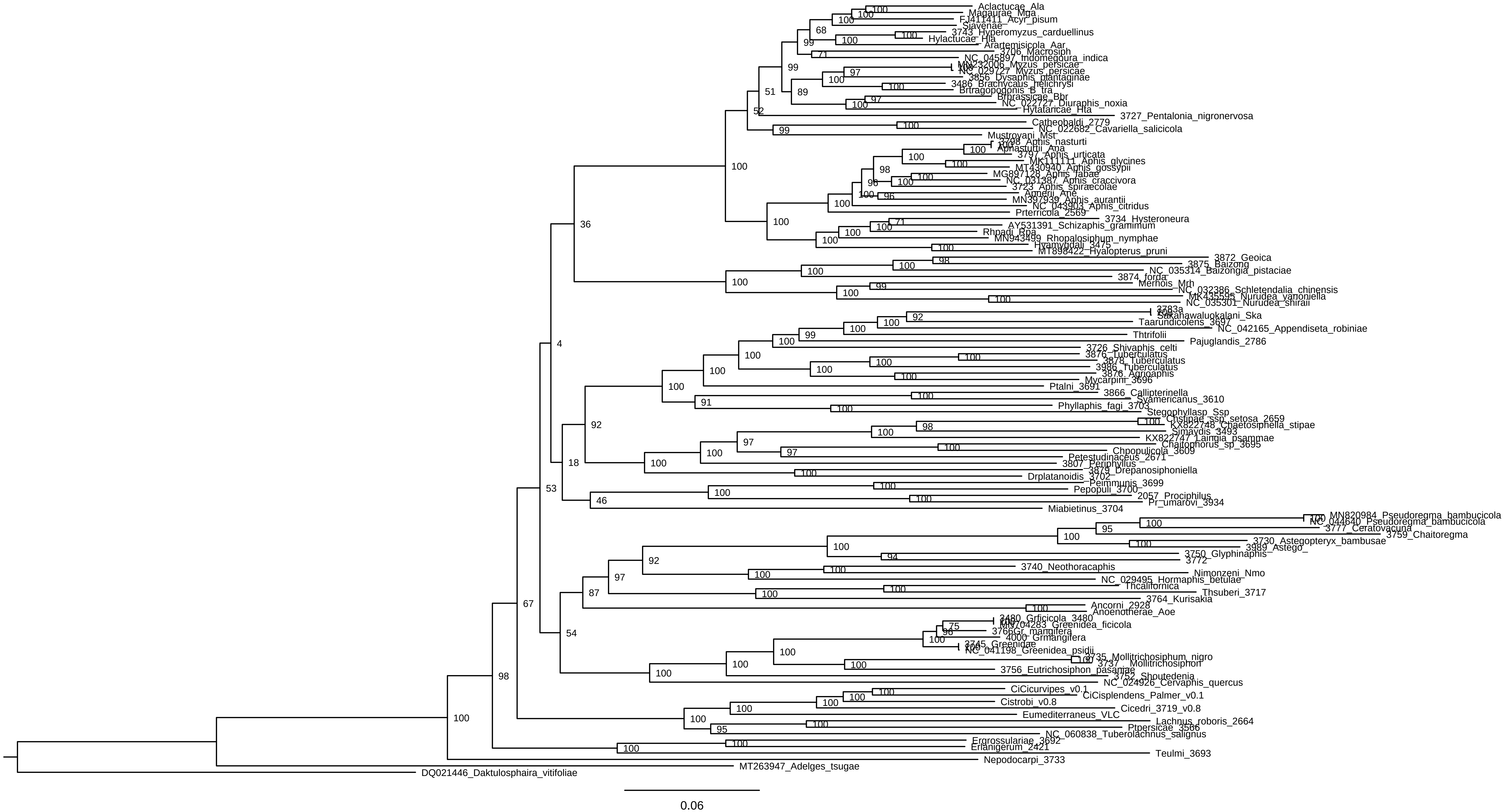

### NucMitoch_Partitioncontree.pdf

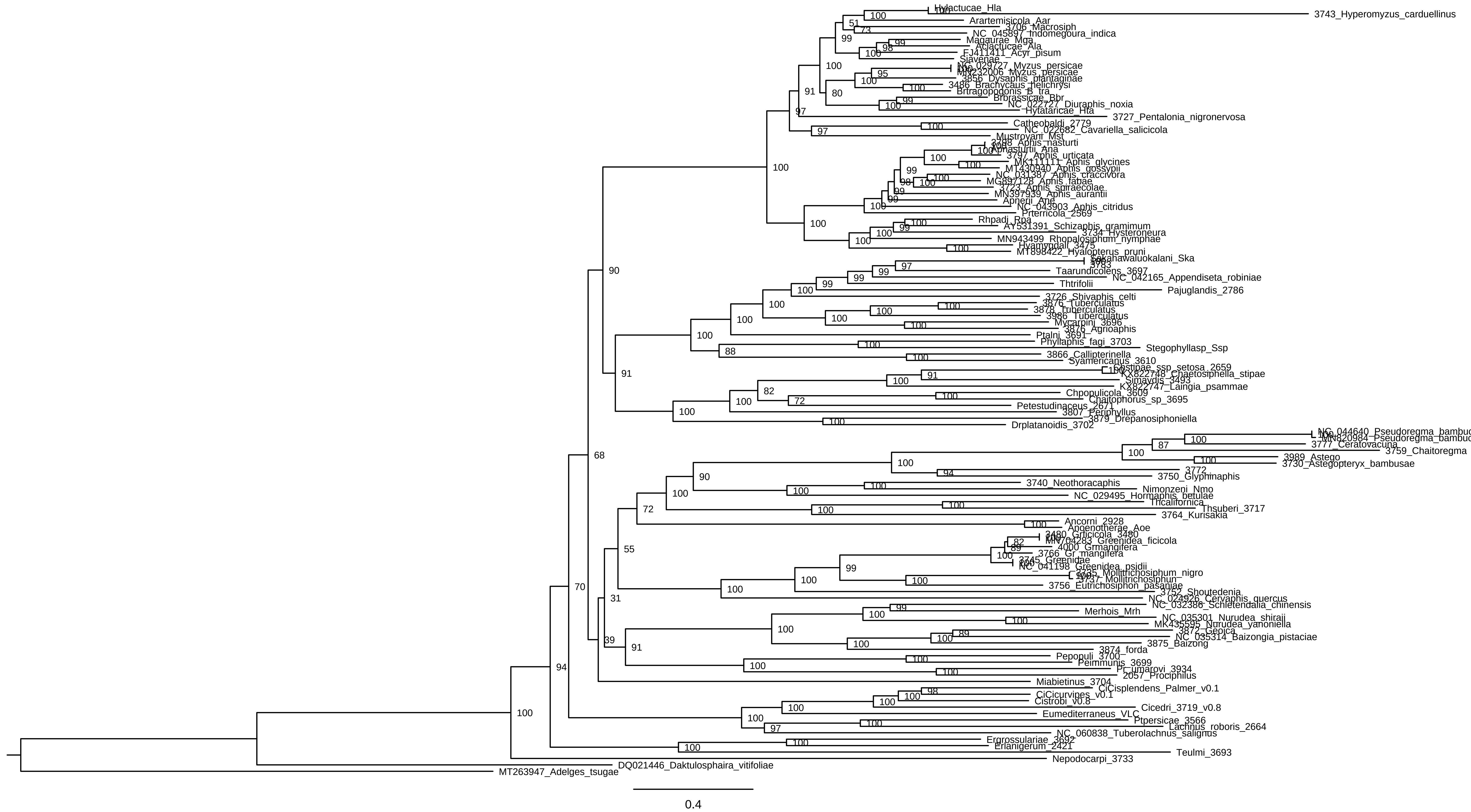

### PartitionsAA_Mitochcontree.pdf

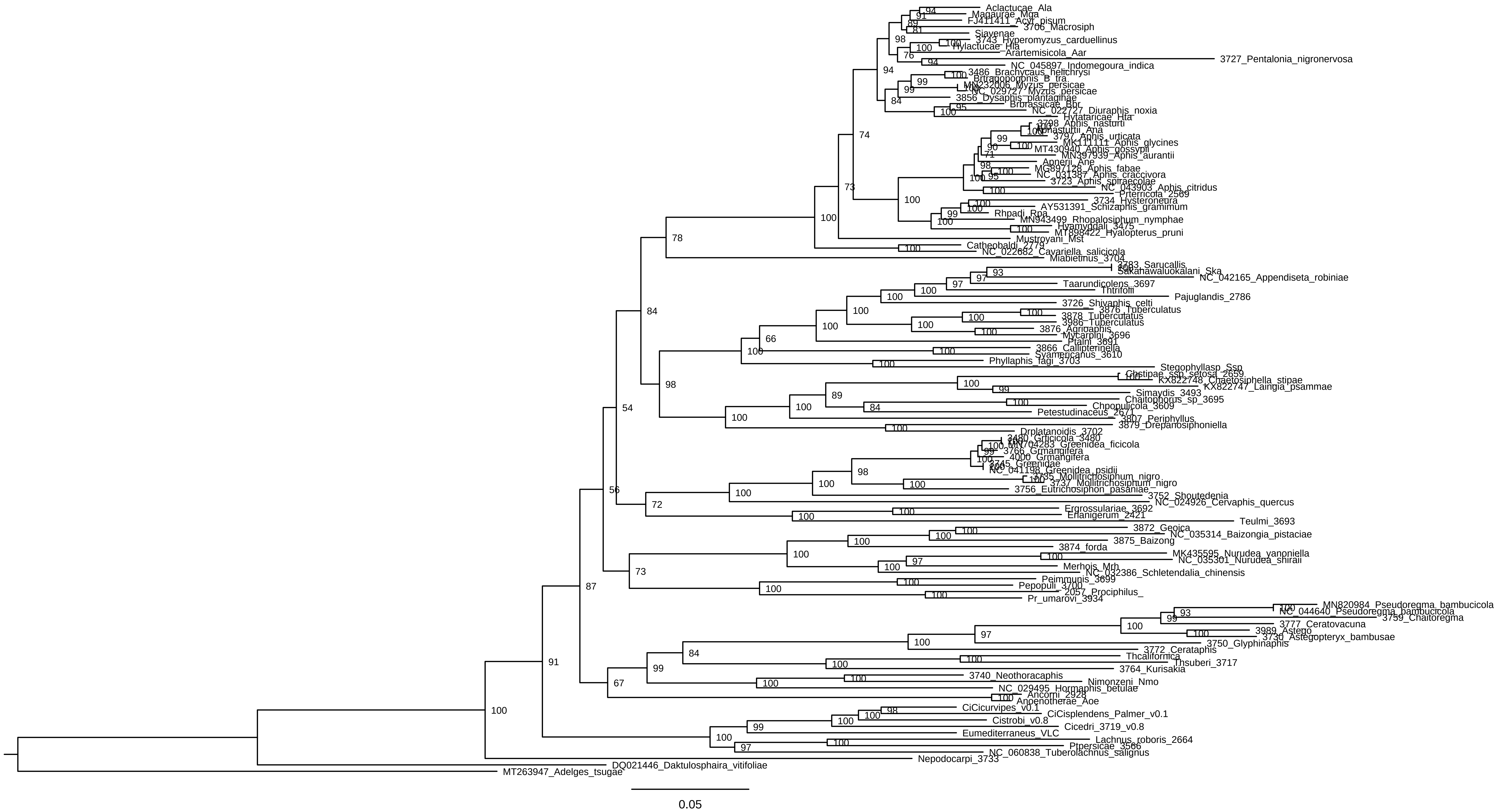

### PhyloBayes_AA_Mitoch.pdf

Seaview PhyloBayes_AA_Mitoch.con.tre Fri Apr 12 09:15:16 2024

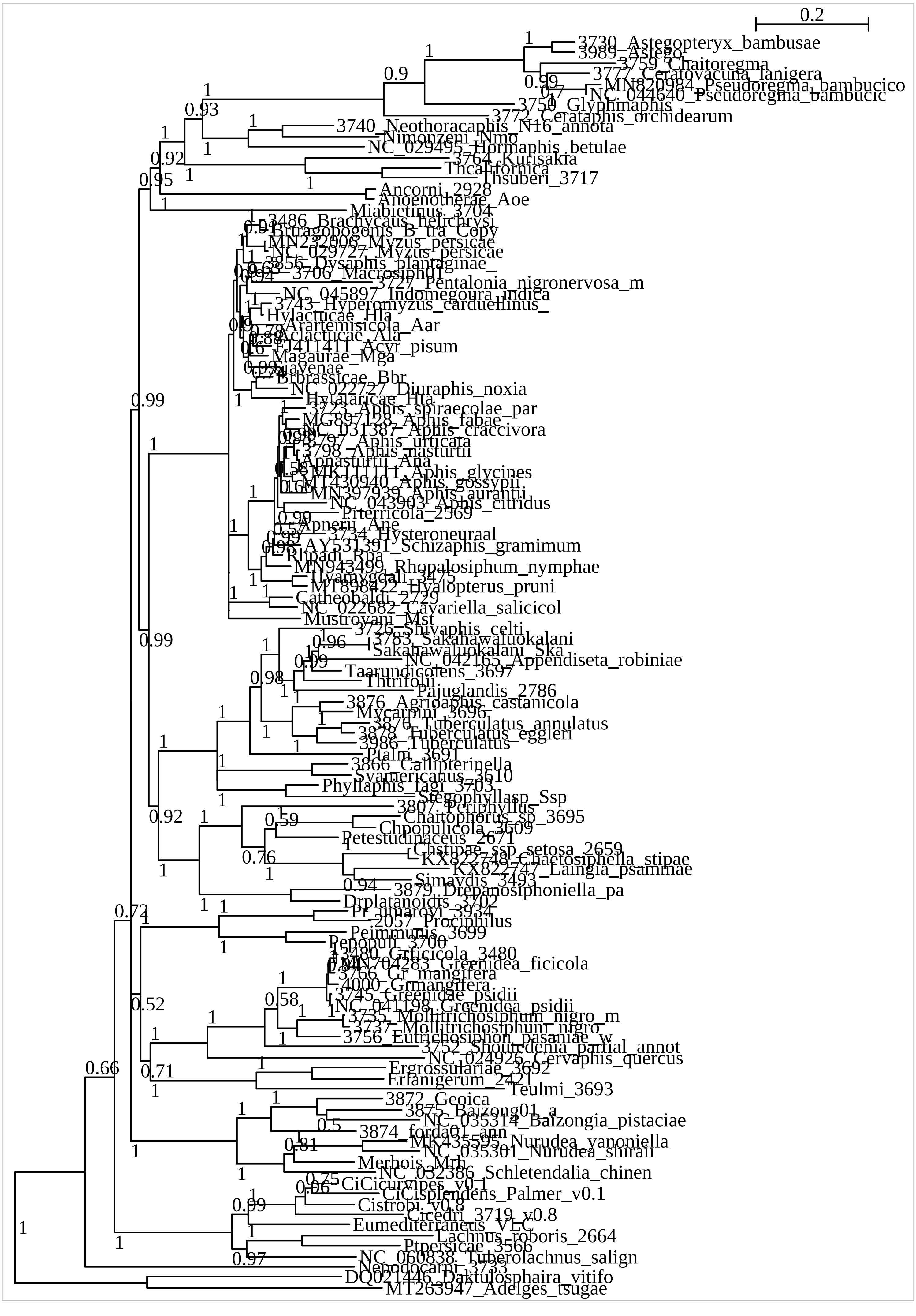

### PhylobayesBuchneraPhylogeny.pdf

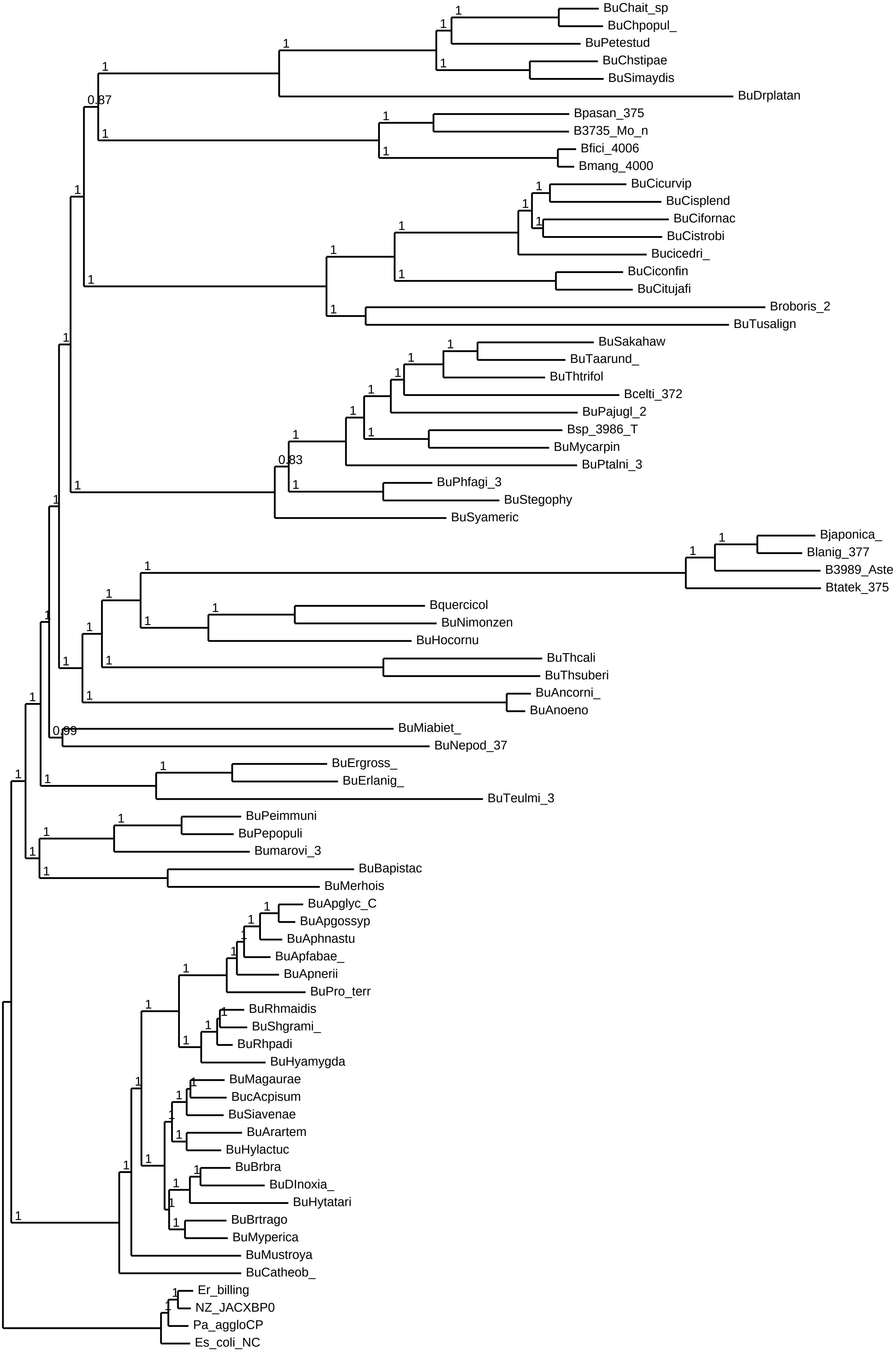
