## Supplementary material for "Discordance between phylogenomic datasets in aphids: who is telling the truth?": TextS1

**Test1:Supplementary material**

**Obtention of mitochondrial genomes through long-range DNA amplification and Illumina sequencin**g

We defined three primer pairs for the amplification of long-range DNA fragments. Those were designed using reference mitochondrial genomes from aphid species from different subfamilies using primer 3 as implemented in Geneious v 11.1.5. We used forward primer LepLR_F 5’-CAACTAATCATAAAGAYATTGG-3’ and reverse primer COIIILR_R TAATRTCAATRAARTGTCAATATCA-3’ to amplify an approximately 4kb long fragment between COI and COIII genes; we used forward primer COIII_LR_F 5’-ATAATYTTATTTATTATTTCWGAA-3’ and reverse primer 16S_LR_R 5’-CAAAGGTAGCRTARTAATTAGTC-3’ to amplify an approximately 8kb long fragment between COIII and 16s rDNA genes, and forward primer 16SrevSens_F 5’-CTTAATTCAACATCGAGGTCGCAA-3’ and reverse primer Lep_LR_R 5’-TCTATWCCAATWGTAAATATAT-3’ to amplify an approximately 6 kb fragment between the 16S rDNA gene and COI gene.

LA Taq HS DNA polymerase kit from TaKaRa (TaKaRa Bio) was used for DNA amplification. For each DNA fragment, for each sample, long range PCR was performed in a total volume of 25 µL containing 2.5 µL of 10x TaKaRa buffer, 0.25 μl Taq polymerase, 2.5 µL of each primer (0.7 μM), 4 μl dNTPs (10 mM), 2.5 µL of Mgcl2 and 9.75 μl nuclease free water. PCR conditions consisted of an initial denaturation step at 94°C for 4 min followed by 35 cycles of denaturation at 98°C for 10s, hybridization at 51°C for 30 s and extension at 60°C for 5 min to 10 minutes depending of fragment length, and a final extension step at 72°C for 10 min. After PCR, 1 µl of the reaction mixture was used for electrophoresis on a 0.4% agarose gel. PCR amplification success is given on in Table SX.

For each DNA sample, we pooled long-range PCR products in a single sample, those were then submitted to Nextera library preparation. Two ng input DNA per sample was used for library preparation using the Nextera XT library preparation kit, and samples were indexed using the Nextera XT index kit (Illumina). After index PCR, samples were purified with 45 μl of Agencourt AMPure XP magnetic beads with a sample to beads ratio of 3:2 (Beckman Coulter), normalized by Qubit quantification and pooled to generate a 4 nM library. To compensate for low diversity libraries, a 12 pM PhiX control spike-in of 2.5% was added. Paired-end sequencing (V2 kits) was done on an Illumina MiSeq.

**Two step PCR protocol: final step of Fluidigm amplification**

A second PCR step was performed to add individual-specific multiplexing tags (called index i5 and index i7, see below), consisting of short 9 bp sample-specific sequences (Martin et al 2019) and the Illumina adapters (called P5 and P7) at the 5' ends of each amplified DNA fragment to the first PCR1. As all the PCR2 products were mixed together (multiplexing) for MiSeq sequencing, dual-indexes made it possible to identify the origin of the sequences and reassign them to each sample (demultiplexing). Each i5 and i7 indexes were used for a unique PCR replicate (i.e. unique dual indexes) to reduce the index-hopping and make sure that libraries were sequence and demultiplex with the highest accuracy (Kircher et al 2012). This second PCR was carried out in a total volume of 11 µL containing 5 µL of Multiplex kit (Qiagen, Germany), 0.7 μM of each primer, and 2 µL of products from the first PCR for each sample. PCR conditions consisted of an initial denaturation step at 95°C for 15 min followed by 8 cycles of denaturation at 95°C for 40 s, hybridization at 55°C for 45 s and extension at 72°C for 2 min, and a final extension step at 72°C for 10 min.

Forward indexed i5 primer PCR2 sequence :

AATGATACGGCGACCACCGAGATCTACAC<i5>TCGTCGGCAGCGTC

Reverse indexed i7 primer PCR2 sequence:

CAAGCAGAAGACGGCATACGAGAT<i7>GTCTCGTGGGCTCGG)
